# Novel apoptosis signal-regulating kinase 1 (ASK1) inhibitor SRT-015: Potential therapeutic for multiple liver diseases

**DOI:** 10.64898/2026.06.30.735673

**Authors:** Kathleen A Elias, Samuel D Brown, Michael Feigh, Neil D. McDonnell, Artur Plonowski

## Abstract

**Background & Aims:** Activation of apoptosis signal-regulating kinase 1 (ASK1), a ubiquitous redox-sensitive kinase, results in inflammation, apoptosis, and fibrosis, key common pathways in human liver disease. SRT-015 is a novel, small molecule inhibitor of ASK1. This study evaluated the in vitro efficacy of SRT-015, compared it to other ASK1 inhibitors, and determined the in vivo efficacy of SRT-015 across multiple acute and chronic liver disease models.

**Methods:** In vitro studies determined the kinase potency and selectivity of SRT-015, and cellular studies were used to demonstrate direct mechanisms of action. The cardiac hERG channel inhibition was assessed and PK determined in rodents and nonhuman primates. In vivo studies evaluated SRT-015 efficacy in rodent models of drug-induced hepatotoxicity (acetaminophen (APAP) overdose), alcohol-associated liver disease (ALD), metabolic-disease associated steatohepatitis (MASH) and cholestatic disease (bile duct ligation, BDL).

**Results:** SRT-015, was demonstrated a selective ASK1 kinase, and SRT-015 treatment directly inhibited fibrosis, apoptosis and inflammation in activated human fibroblasts, hepatocytes and PBMCs, respectively without safety signals or hERG inhibition. Other ASK1 inhibitors had safety concerns or limited functional activity. Liver and kidney selective PK were observed for SRT-015 in all species evaluated. In vivo, SRT-015 treatment was efficacious in the acute mouse APAP overdose and ALD model significantly (P<0.05) decreasing serum ALT. Using a therapeutic diet-induced obesity (DIO)-MASH model with biopsy-verified fibrosis, SRT-015 treatment significantly (P<0.05) inhibited DIO-induced liver enzymes, hepatomegaly, fibrosis, inflammation, and apoptosis independent of body weight loss whereas treatment with selonsertib was ineffective. In a rat cholestatic model, SRT-015 treatment significantly (P<0.05) decreased fibrosis and stellate cell activation.

**Conclusions:** These findings support SRT-015 as a potential therapeutic for human liver diseases of any etiology.

## Introduction

The global burden of chronic liver disease is rising with cirrhosis, or end-stage liver disease, being the 11th leading cause of death and the 14^th^ leading cause of morbidity worldwide (1). Hepatic fibrosis is the common pathological mechanism that leads to cirrhosis and can occur following chronic liver injury from toxins (alcohol), infections (HAV and HBV), cholestasis, and metabolic-disease associated steatohepatitis (MASH, formally known as NASH; 2).

Regardless of etiology, key pathology in liver disease progression includes inflammation, fibrosis and apoptosis driven by crosstalk between the different cell types, including hepatocytes, immune cells and hepatic stellate cells (2–5).

Illustrating the heterogeneity of these diseases, specific liver cell types are activated by different stress factors (6). For example, in MASH hepatocytes are primarily affected by lipotoxicity, which is further dependent on the lipid type (7). Kupffer cells, the resident macrophages in the liver are activated by hepatocyte death and by the endotoxin lipopolysaccharide (LPS) from the portal blood via the gut-liver axis (8) to initiate the inflammatory response (9). This induces macrophages and neutrophils to migrate into the liver and release more pro-inflammatory cytokines that, in combination with stressed or injured hepatocytes, in turn activate the stellate cells to produce matrix proteins and initiate the fibrotic response (10). The “multiple-hits” hypothesis suggests that in addition to fat accumulation, oxidative and endoplasmic reticulum (ER) stress also drives liver inflammation and fibrosis (11–13).

Extensive fibrosis results in cirrhosis, a common endpoint in MASH, ALD and cholangiopathies. A key common pathway in all these liver diseases is the activation of ASK1 (14–17).

ASK1 (MAP3K5) is a ubiquitous redox-sensitive kinase that is activated by pathological stimuli including reactive oxidative stress (ROS), lipotoxicity, cytokines and ER stress (16). Inactive ASK1 is localized in the cytoplasm and mitochondria, and is normally bound and repressed by antioxidant proteins, including thioredoxin 1 in the cytosol and thioredoxin 2 in the mitochondria (18). Under oxidizing conditions ASK1 is activated, leading to fibrosis, inflammation, and apoptosis as described in multiple systems (19–21).

In the liver, diverse stress stimuli including lipotoxicity, alcohol, cytokines, cholestatic and hepatotoxic agents, induce the formation of ROS resulting in the oxidation and dissociation of thioredoxin from ASK1, leading to ASK1 activation. In turn, activated ASK1 induces phosphorylation and activation of c-Jun N-terminal kinase (JNK) and p38 MAP downstream kinase cascades (22). ASK1 is activated in hepatocytes, Kupffer cells and stellate cells in the liver, resulting in hepatocyte cell death, inflammation and fibrosis, respectively.

Inhibition of activated ASK1 is a well-validated preclinical target for multiple disorders including liver, kidney, cardiovascular, lung and CNS diseases (21,23). Despite extensive preclinical evidence of the therapeutic potential of ASK1 inhibition, selonsertib, the first ASK1 inhibitor to enter clinical trials, was not efficacious in clinical trials for MASH (24) or ALD (25) likely due to recognized off-target liabilities (26–30).

SRT-015 is a novel small molecule inhibitor of the activated ASK1 kinase identified in a drug development program at Seal Rock Therapeutics without the off-target liabilities of selonsertib. SRT-015 occupies the ATP-binding pocket of ASK1, preventing its auto-phosphorylation and the activation of downstream signaling via JNK and p38 mitogen-activated protein kinases. SRT-015 is being developed as a possible treatment for multiple liver diseases including MASH, ALD and cholangiopathies. Here we report SRT-015 in vitro and in vivo pharmacology results including the demonstration of direct inhibition of ASK1-induced fibrosis, inflammation and apoptosis in the specific cell types responsible for the pathology. No safety concerns were observed with SRT-015 treatment, in contrast to the other ASK1 inhibitors including selonsertib. SRT-015 has liver-selective pharmacokinetics (PK) with preclinical efficacy in liver models including acute ALD and APAP hepatotoxicity models. Using a rat cholestatic model, SRT-015 significantly decreased fibrosis and stellate cell activation. In a chronic, therapeutic DIO-MASH model with biopsy-verified fibrosis, SRT-015 treatment demonstrated efficacy by directly inhibiting established fibrosis, inflammation and apoptosis. These findings demonstrate that SRT-015 is a potent and selective ASK1 inhibitor with a pleotropic mechanism that may be a beneficial therapeutic agent for treating human liver diseases.

## Materials and Methods

### Compounds

SRT-015 (31), Takeda 19 (32), Pfizer 18 and Pfizer 38 (33) were synthesized by SYNthesis Med Chem (Parkville, VIC, AU). Selonsertib (GS-4997) was obtained from Chemietek (Indianapolis, IN, USA), GS-444217 (ProbeChem, Shanghai, China) and the JNK inhibitor SP600125 and p38 inhibitor Losmapimod from Selleckchem (Houston, TX, USA).

### Biochemical assessments

#### Enzymatic

SRT-015, other ASK1 inhibitors and reference compounds were measured in a 10-pt dose response format and the half-maximal inhibitory concentration (IC_50_) was determined by the inhibition of recombinant human ASK1 enzyme (ADP-GloTM Kinase assay + ASK1 kinase; Promega, Madison, WI, USA).

#### Kinome

Kinome selectivity of SRT-015 (1μM) was determined using the KINOMEscan binding assay (DiscoverX, Hayward, CA, USA) against 489 kinases. The top kinase inhibitors (0.1% control or > 90% kinase inhibition) were identified and further assessed using 11-point Kd dose response curves in duplicate (KdElect; DiscoverX, Hayward, CA, USA) or IC_50_ assays (Reaction Biology, Malvern, PA, USA).

#### Off-target safety assessments

SRT-015 (10 μM) was evaluated using SafetyScreen87^TM^, a primary molecular target panel consisting of 13 enzymatic and 74 binding assays designed to identify potential off-target liabilities (Eurofins Panlabs Discovery Services, Taipei, Taiwan) and automated clamp assays of the human ether-a-go-go-related gene (hERG) channel (WuXi Biologics, Shanghai, China) with SRT-015 and reference compounds.

### Cell Culture and Mechanism Assays

#### Cells

HepG2 cells and primary human dermal fibroblasts were obtained from ATCC (Manassas, VA, USA) and primary human cryopreserved normal peripheral blood mononuclear cells (PBMC) were obtained from ALLCells (Alameda, CA, USA).

#### Study design

Depending on the cell type, cells were challenged using cell appropriate stress protocols to induce ROS and compounds were evaluated in a dose-dependent manner to determine the inhibition of TNF, p-p38, caspase3/7 or alpha-smooth muscle actin (α-SMA), as described below.

#### Anti-inflammatory MOA and target engagement

To evaluate the TLR4-ASK1-p38 pathway (34,35) human PBMCs were plated in 96-well TC plates in RPMI media with heat-inactivated FBS (ThermoFisher, South San Francisco CA, USA). PBMCs were challenged with 100 ng/ml LPS (Sigma, Livonia, Michigan, USA) for 20 min to activate ASK1 and determine p-p38 levels (five donors; HTRF P-p38 Kit; CisBio, Codolet, France) and 6 h for TNF levels using a sandwich ELISA (nine donors, TNF; BD OptEIA, Franklin Lakes, NJ). Compounds were added 1 h prior to LPS challenge and tested in a 10pt dose response format (0.0015 – 30μM in 0.3% DMSO) to determine percent inhibition values plotted against compound concentration. Dose-response curves were fitted by a 4-parameter sigmoid inhibition model using the XLFit® software (IDBS, Oakland, CA, USA) to determine IC_50_ values.

#### Anti-apoptosis MOA

HepG2 cells were plated in two 96-well plates at a density of 30,000cells/well in 100uL of EMEM supplemented with 10% FBS and incubated for 48h at 37°C in 5%CO2. After 48h of incubation, the medium was replaced with 50μL of phenol-free DMEM supplemented with 10%FBS and 4.5g/L glucose. The compounds were serially diluted and added to the cells (0.94-30μM). The final DMSO concentration was equalized at 0.3% in all experimental and control samples. After 1h preincubation with compounds, the cells were stimulated with 1mM H_2_O_2_ for 20h to induce apoptosis. After stimulation, one of the assay plates was used to determine Caspase3/7 activity using Caspase3/7-Glo® detection reagent (Promega, Madison, WI, USA) pre-diluted 1:1 in DMEM medium without phenol red and the second plate was used to determine overall cell viability using Cell-Titer Glo® reagent. Luminescence was measured on a BioTek Synergy Neo2 instrument (Agilent Technologies, Santa Clara CA, USA). Caspase 3/7 activity was normalized to the cell viability at each concentration point. The assay window was determined in each experiment by comparing the basal Caspase3/7 activity in untreated cells to that in cells treated with H_2_O_2_, in the absence of compound. The percent decrease in apoptosis compared to H_2_O_2_ controls was used only when cells exhibited 95% or higher viability as other cell killing mechanisms such as necrosis, may occur in stressed cells.

#### Antifibrotic MOA

Human fibroblasts were maintained in basal fibroblast growth medium supplemented with 2% FBS serum kit (ATCC, Manassas, VA, USA) in 96 well TC plate at a density of 4000cells/well and incubated for 72h at 37 °C in 5%CO2 for 72 h. The medium was replaced with DMEM supplemented with 1% FBS, and the cells were serum-starved for 48h. After 48 h, serially diluted compounds (0.37 –30 μM) in 1% FBS DMEM with DMSO concentration 0.3% in all test wells was added. After 1 h preincubation with compounds, 0.3ng/mL TGF (R&D Systems, Minneapolis, MN, USA) was added to induce differentiation of fibroblasts into myofibroblasts. After 48 h of incubation, α-SMA was determined using the reagents and instructions per the In-cell ELISA kit (ThermoFisher, South San Francisco, CA, USA), an anti-α-SMA antibody (1A4, Abcam, Fremont, CA, USA) and SpectraMax plate reader (Molecular Devices, San Jose CA, USA). The calculated percent inhibition values were plotted against the compound concentration. To determine the IC_50_ values, dose-response curves were fitted by a 4-parameter sigmoid inhibition model using the XLFit® software (IDBS, Oakland, CA, USA).

### Cytotoxicity

To determine the cytotoxic effects of compounds in unstimulated cells (no stress protocol with H_2_O_2_ or TGF), human dermal fibroblasts and HepG2 cells were plated in DMEM or EMEM media and allowed to undergo three population doublings and cytotoxicity was measured by Cell-Titer Glo (Promega Madison, WI, USA).

#### Animal studies

Animal experiments were conducted at contract research organizations in facilities accredited by the Association for Assessment and Accreditation of Laboratory Animal Care International (AAALAC). All experimental protocols were approved by the Institutional Animal Care and Use Committees to ensure compliance with the Guide for the Care and Use of Laboratory Animals (NIH) or other applicable local regulations.

#### Pharmacokinetics

Pharmacokinetic parameters were determined in plasma samples obtained serially after a single oral (PO, 5 mg/kg) administration of SRT-015 in male Wistar rats, male cynomolgus monkeys, and male C57BL/6 mice (PO, 10 mg/kg). The liver and other tissue distributions of SRT-015 were also determined.

#### Bioanalysis

All blood and tissue samples were processed for analysis by protein precipitation using acetonitrile and analyzed using a fit-for-purpose LC-MS/MS method (LLOQ = 2.49 ng/mL).

### Therapeutic acute acetaminophen (APAP) hepatotoxicity model

#### Animals

Male C57BL/6 wild-type mice (20-22 g) were acclimated for at least three days in a temperature-controlled (22 ± 3 ^0^C), 12-h light/dark room.

#### Study design

In the APAP therapeutic mouse model, APAP (300 mg/kg; i.p.) was administered and 1 h later SRT-015 (0.3-10 mg/kg), or vehicle (5% N-methylpyrrolidone (NMP) + 10% Solutol + 55% PEG400 + 30% water) was administered by oral gavage (n = 6/group). Six hours after APAP administration, ALT levels were measured in terminal plasma. In a separate experiment, livers from treated mice were isolated at 6 h post-treatment with APAP and equal amounts of pooled liver protein (50 μg) from each group were subjected to SDS-PAGE and western blotting for JNK, p-JNK, ASK1, p-ASK1, p38 and p-p38; the same blots were stripped and re-probed with anti-α-SMA Ab.

### Pyrazole LPS model of acute ALD

#### Animals

Male C57BL/6 wild-type mice (20-242 g) were acclimated for at least three days in a temperature-controlled (22 ± 3 ^0^C), 12-h light/dark room.

#### Study design

Pyrazole was administered on two consecutive days to induce CYP2E1 prior to SRT-015 (BID) and LPS (single dose) administration. LPS administration emulates the inflammation caused by gut bacteria penetrating the intestinal wall following binge drinking. Mice were divided into five groups: saline (10 mL/kg; excipient for pyrazole and LPS), pyrazole (150 mg/kg) and LPS (4 mg/kg) administered by i.p. injection. SRT-015 (1,3,10 mg/kg) or vehicle (10 mL/kg) was administered by oral gavage BID. Mice were terminated 24 h after LPS administration and serum and plasma were collected for ALT and bioanalysis.

### Therapeutic DIO-MASH mouse model with fibrosis

#### Animals

Male wild-type C57BL/6J mice were fed a DIO-MASH diet high in fat (40% total fat kcal/ 18% trans-fat), fructose (20%) and cholesterol (2%) for 38 weeks using a previously described protocol (39). One week before compound treatment initiation, all mice underwent liver biopsy for confirmation and stratification of liver steatosis and fibrosis and only biopsy-confirmed mice with steatosis score >2 and fibrosis stage >1 were used for the study.

#### Study design

After 38 weeks on DIO-MASH diet, mice with biopsy-confirmed steatosis and fibrosis were kept on the DIO-MASH diet and stratified into groups (n = 11-12) including DIO-MASH diet control, SRT-015 treatment (0.3% or 0.5% in DIO-MASH diet), or selonsertib (0.1% in DIO-MASH diet) for an additional 12 weeks. Selonsertib dosing concentration was designed to match the clinical trial plasma concentrations and match SRT-015 liver exposures. Age-matched control mice (Lean Chow) were maintained on a rodent chow diet for the duration of the study. At 50 weeks, mice were evaluated by histological, biochemical, and RNAseq methods as described previously (36). The SRT-015 0.3% group did not undergo RNAseq analysis.

#### Target Engagement p-p38 IHC DIO-MASH livers

Liver sections from the DIO-MASH mouse study were additionally assessed by IHC for p-p38 (Cell Signaling, Danvers, MA, USA) activity in hepatocytes and non-hepatocytes (immune and stellate cells) populations using an AI-powered digital assay pathology platform for cellular segmentation and analysis (Reveal Biosciences, San Diego, CA, USA).

### Bile Duct ligation model

#### Animals

Male Sprague Dawley rats (7 - 8 weeks) were randomized into four treatment groups: sham control (n = 4), BDL + vehicle (n = 10), BDL + 10 mg/kg SRT-015 (n = 10) or BDL + 50 mg/kg SRT-015 (n = 10). BDL surgery included ligation of the common bile duct below the junction of the hepatic ducts and above the pancreatic duct entrance. Sham animals underwent laparotomy without ligation.

#### Study design

SRT-015 or vehicle was dosed po (80:20 Capryol90: Labrafil M 1944 CS, % v/v) starting at Day1, 1 h before surgery, then daily for 14 days. On terminal Day 14, 2 h after the last dose, whole blood was collected for blood chemistry evaluation. Livers were evaluated for SRT-015 concentrations, fibrosis by biochemical hydroxyproline (HP) content, picrosirius red staining (PSR), and stellate cell activation (α-smooth muscle actin staining) by histomorphometry (Aperio ImageScope analysis software Leica Biosystems, Deer Park, USA).

#### Statistical analysis

For all studies the values are expressed as mean ± SEM and statistical analysis was performed using One-way ANOVA with Dunnett’s post-hoc test or Tukey’s multiple comparisons test using GraphPad Prism software except where noted.

## Results

### In Vitro

#### SRT-015 kinase inhibition, selectivity, and off-target assessments

##### SRT-015 potent ASK1 inhibitor

SRT-015 inhibition (IC_50_) of the ASK1 ATP binding domain was determined to be 24.4 +/- 1.9 nM (n = 23) in enzymatic assays using a 10-point curve from 10 different SRT-015 research and production lots (Fig. 1A.)

**Fig. 1.**
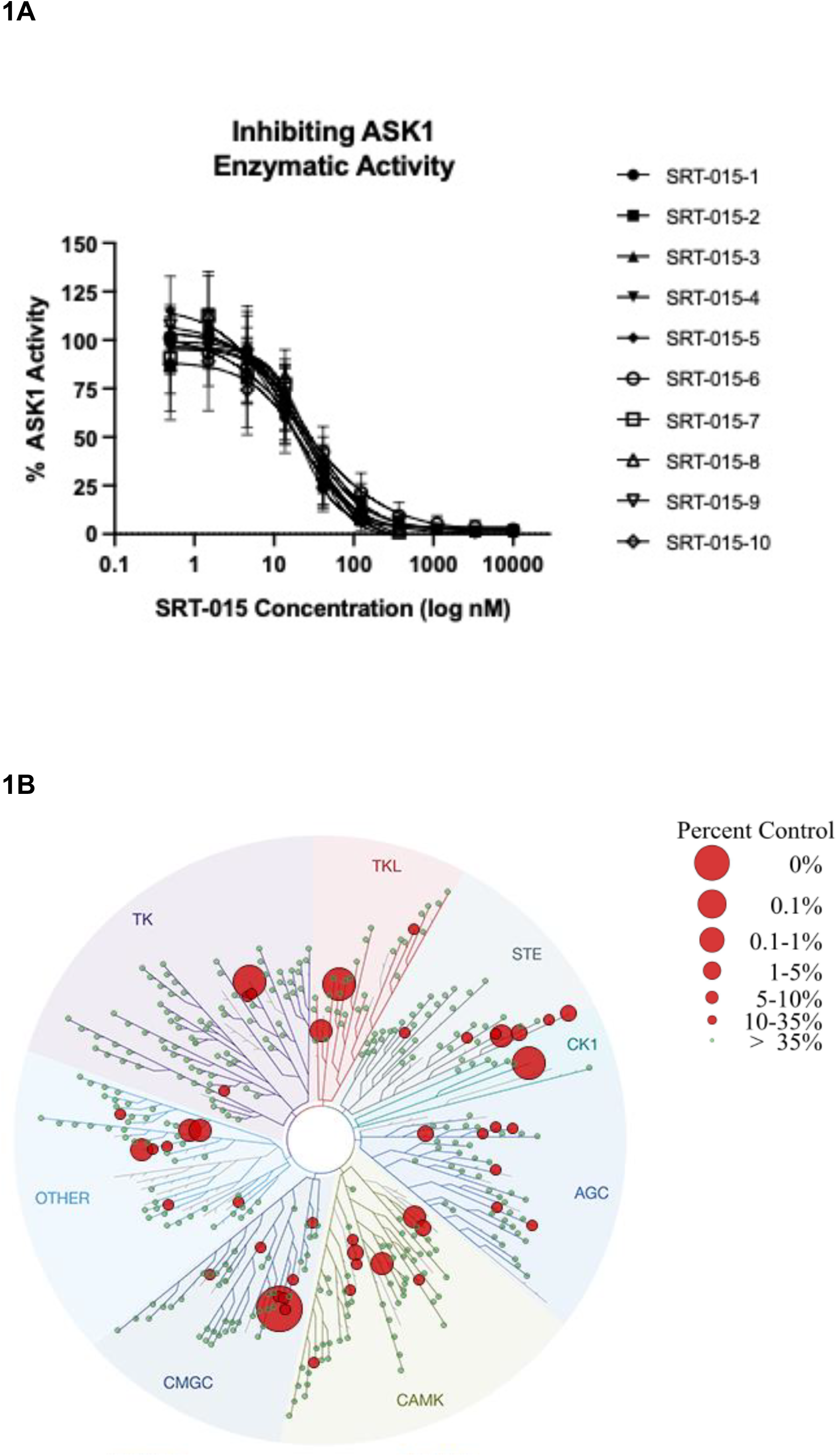
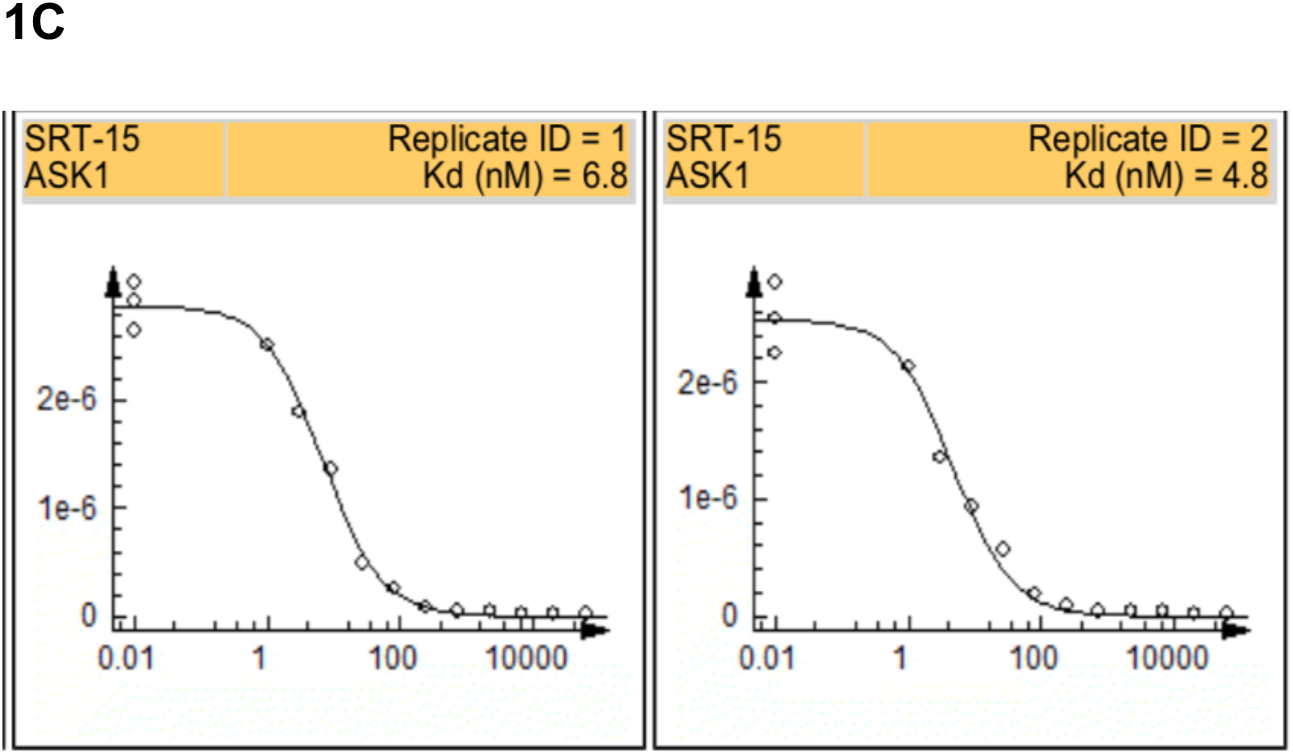
SRT-015 ASK1 inhibition, kinase profile, and Kd dose response. (A) SRT-015 dose-response curves demonstrating increasing inhibition of ASK1 in 10-pt enzymatic assay using 10 different SRT-015 research and production lots (B) SRT-015 evaluated at 1 μM in a full kinome scan (489 kinases KinomeScan) resulting in the inhibition at 95% or greater of 11 kinases. All 11 kinases were evaluated in full Kd dose response with four kinases identified as the primary kinases inhibited by SRT-015 (C) SRT-015 11-point Kd dose-response binding curves to ASK1 in duplicate.

##### SRT-015 selective kinase profile

To determine kinase selectivity, SRT-015 was evaluated at 1 μM in a full kinome scan (489 kinases) resulting in the inhibition of 11 non-mutated kinases by 95% or greater (Fig. 1B). The follow-up full Kd or IC_50_ determinations showed that ASK1, DRAK2, ASK3 and LRRK2 were the key kinases inhibited by SRT-015 with an ASK1/kinase ratio of <10. SRT-015 was the most potently bound to ASK1 averaging 5.8 nM (Fig. 1C). ASK1 and ASK3 are additional targets of interest while inhibition of DRAK2 (STK17B) may be beneficial for chronic liver diseases due to its critical role in HCC progression (37).

##### SRT-015 does not bind off-target receptor panel or hERG channel

No off-target binding (≥ 50% inhibition at 10 μM) was observed with SRT-015 using the SafetyScreen87^TM^, a primary molecular target panel. This panel includes molecular targets that are known to be involved in adverse clinical drug reactions. Automated clamp assays of the human hERG channel SRT-015 demonstrated 10% inhibition of this cardiac channel indicating that cardiac QT prolongation is unlikely to occur following SRT-015 treatment. No change with SRT-015 on cardiac QT prolongation was observed in Phase 1 results with SRT-015 (38). Table 1 shows SRT-015 and other ASK1 inhibitors and reference standards hERG channel inhibition.

**Table 1.** Responses of ASK1 inhibitors and reference standards in ASK1 enzymatic, hERG, MOA and cytotoxicity assays. SRT-015 is the only compound demonstrating all the required compound phenotypes including MOA responses (anti-inflammatory, antifibrotic, anti-apoptotic), acute p-p38 inhibition demonstrating target engagement with no hERG inhibition or cytotoxicity. IC_50_ presented as Mean ± SEM; Experimental n in parathesis. ND = not determined.

| Required Phenotype | ASK1 inhibition | <20% hERG inhibition | Downstream Pathway Acute Inhibition (Target Engagement) | Anti-inflammatory MOA | Antifibrotic MOA | Anti-apoptotic MOA | Lack cytotoxicity (human primary cells) | Lack cytotoxicity (human cell line) |
| --- | --- | --- | --- | --- | --- | --- | --- | --- |
| Compound | ASK1 Enzymatic IC<sub>50</sub> (nM) | hERG channel (@10 μM) | hPBMC LPS-induced p-p38 inhibition IC<sub>50</sub> (μM) | hPBMC LPS-induced TNF inhibition IC<sub>50</sub> (μM) | hFibroblasts TGF-induced α-SMA inhibition IC<sub>50</sub> (μM) | HepG2 H<sub>2</sub>O<sub>2</sub>-induced apoptosis inhibition @ 7.5 μM | Fibroblast Cytotoxicity IC<sub>50</sub> (μM) | HepG2 Cytotoxicity (72 h) IC<sub>50</sub> (μM) |
| SRT-015 | 24.4 | 10% | 2.8 +/- 0.4 (5) | 9.1 +/- 1.4 (9) | 13.7 +/- 2.7 (4) | 34% (2) | >30 (3) | >30 (3) |
| Selonsertib | 8.4 | 39% | 28.7 +/- 1.3 (8) | 6.1 +/- 0.6 (11) | 23.6 +/- 2.0 (5) | none (3) | 20.8 +/- 2.6 (3) | 20.6 +/- 2.8 (4) |
| GS-444217 | 23.1 | 80% | 5.1 +/- 1.6 (4) | 5.6 +/- 0.8 (4) | 20.6 (2) | none (2) | >30 (2) | 26.2 (2) |
| Takeda 19 | 15.7 | 45% | 0.4 (1) | 1.1 (2) | 1.2 (2) | none (1) | 7.3 +/- 2.9 (3) | 23.1 (2) |
| Pfizer 18 | 5.8 | ND | >30 (2) | 13.8 (2) | >30 (2) | 28% (1) | >30 (2) | >30 (1) |
| Pfizer 38 | 9.6 | 49% | 1.49 (2) | 1.9 (2) | 6.4 (2) | 12% (1) | >30 (2) | >30 (1) |
| p38 inhibitor Losmapimod | >10,000 | ND | 0.002 +/- .001 (3) | 0.19 +/- 0.1 (3) | 30 (1) | 28% (1) | >30 (1) | >30 (1) |
| JNK inhibitor SP600125 | 397 | ND | >30 (1) | 7.8 (2) | 6.4 (2) | 67% (1) | 8.5 (2) | 29.9 +/- 0.1 (3) |

### Cell Culture and Mechanism Assays

ASK1 activation results in inflammation, apoptosis and fibrosis both in vitro and in vivo (21). To demonstrate the direct inhibition of these cellular pathways by SRT-015, we used human primary cells and human cell lines challenged with ROS or ROS-inducing factors to activate ASK1.

#### SRT-015 anti-inflammatory MOA

SRT-015 inhibited ROS-induced cytokine release in human PBMCs first challenged with 100 ng/mL LPS to activate ASK1 and induce downstream p-p38 and TNF secretion. Incubation with SRT-015 (0.001 – 30 μM) dose-dependently inhibited p-p38 and TNF levels (Table1, sFig. 1A) resulting in an IC_50_ of 2.8 ± 0.4 (n = 5) and 9.1 ± 1.4 (n =9), respectively (μM; Mean ± SEM) demonstrating SRT-015 target engagement and inhibition of cytokine pathways.

#### SRT-015 antifibrotic MOA

Following liver injury, hepatic stellate cells undergo an activation response characterized by the accumulation of excessive extracellular matrix secreted by the activated myofibroblasts. The myofibroblast marker of fibrogenic cell activation, α-SMA, was induced in human fibroblasts cells by TGFβ and inhibited by SRT-015 in a dose-responsive manner (Table 1, sFig.1B) with an IC_50_ of 13.7 ± 2.7 μM (Mean ± SEM, n =4).

#### SRT-015 anti-apoptotic MOA

HepG2 cells, a human liver cell line, were challenged with H_2_O_2_ to result in ROS-induced apoptosis (caspase3/7) at 20 h. Cell viability was determined in parallel and caspase activity normalized to cell viability at each concentration. Caspase 3/7 values were used only when the cell viability was >95%. SRT-015 treatment decreased caspase3/7 in two independent experiments with no significant change in the viability of H_2_O_2_-stressed cells, with a maximum inhibition of H_2_O_2_-induced apoptosis of 34% at 7.5 μM (Table 1, sFig. 1C).

#### SRT-015 lacks off-target cellular proliferation or cytotoxicity

In cells without ROS-inducing stimulus added, ASK1 is not activated and there should be no effects on proliferation or cytotoxicity with SRT-015 treatment unless there is an off-target effect. No changes in viability were observed (IC_50_ >30 μM, n = 3) in HepG2 cells or in primary fibroblasts (Table 1, sFig. 1D and E).

#### In vitro comparisons with other ASK1 inhibitors and reference compounds

Literature identified ASK1 inhibitor compounds (Table 1) demonstrated enzymatic activity against the ASK1 kinase but varied by limited acute p-p38 inhibition, lacking anti-ASK1 MOA effects or displaying off-target cytotoxicity. Only SRT-015 treatment met all the required criteria for cellular mechanisms and safety. Significant levels (>20% at 10 μM) of hERG inhibition were identified in all other ASK1 inhibitor compounds evaluated. All compounds inhibited TNF release at 6 h but some did not via the TLR4-ASK1-p38 cascade indicating a different cellular mechanism. Inhibition of apoptosis was not observed with selonsertib, GS-444217 or Takeda 19 because of their dominant cytotoxic effect. Cytotoxicity was also observed with Takeda 19 and selonsertib in primary fibroblasts and in HepG2 cells when no stress induction was present.

### Pharmacokinetics

SRT-015 is an orally bioavailable compound with a rapid preferential distribution to liver tissue and low levels in peripheral circulation in mice, rats, and cynomolgus monkeys. The time course of SRT-015 distribution to the liver, kidney, heart and plasma was investigated in mice after oral administration of 10 mg/kg (Fig. 2A and B). SRT-015 was distributed rapidly to liver tissue, and to a lesser extent to the kidney and heart tissues. The tissue-to-plasma ratio of SRT-015 determined at the time of maximal concentration was 69 in liver, 6.3 in kidneys, and 3.7 in the heart. Over time, the tissue-to-plasma ratios remained approximately constant, indicating a lack of accumulation in the organs analyzed in this study.

**Fig. 2.**
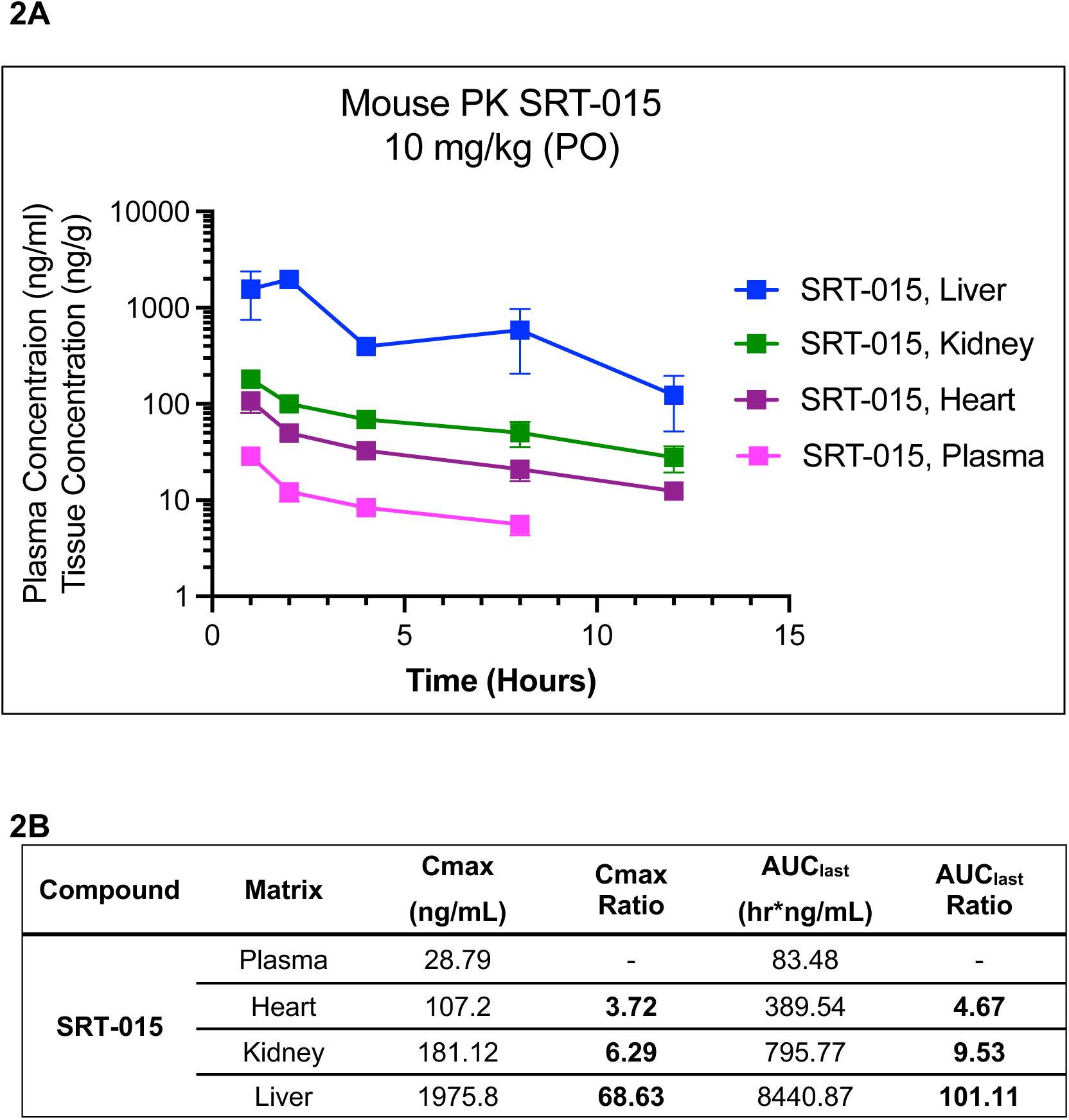
SRT-015 PO PK showing liver-selective exposure. Time course of SRT-015 plasma, liver, heart and kidney concentrations after a single oral dose (10 mg/kg, po) in the mouse (A) and PK parameters (B).

The distribution of SRT-015 to the liver and other organs was further investigated in rats and monkeys (sTable 2). At 1 h after oral administration of 5 mg/kg to rats, the liver/plasma ratio of SRT-015 was 60.4. In monkeys, liver concentrations were determined in three liver lobes 1 h after oral administration and showed uniform distribution within the liver with corresponding liver/plasma ratios at these distinct anatomical sites of 52, 62, and 62, respectively. Similar to the distribution in mice, SRT-015 in monkeys was also substantially distributed in the kidneys (kidney/plasma ratio of 30.9) and to a lesser extent in the heart (heart/plasma ratio of 5.89). As determined in rats, the SRT-015 concentrations in the brain were below the lower limit of quantitation (<15.15 ng/g) indicating a lack of brain penetration.

### Therapeutic acute APAP liver injury model

As shown in Fig. 3A and 3B a single i.p. injection of APAP (300 mg/kg) markedly increased serum ALT and liver p-JNK, with modest increases in p-ASK1 and p-p38 compared to the vehicle-treated mice. Treatment with SRT-015 dose dependently decreased (P<0.001) serum ALT and reduced the phosphorylation of ASK1, JNK and p38 demonstrating the efficacy and target engagement of SRT-015. The minimum efficacious dose (MED) of SRT-015 in this model was 0.3 mg/kg.

**Fig. 3.**
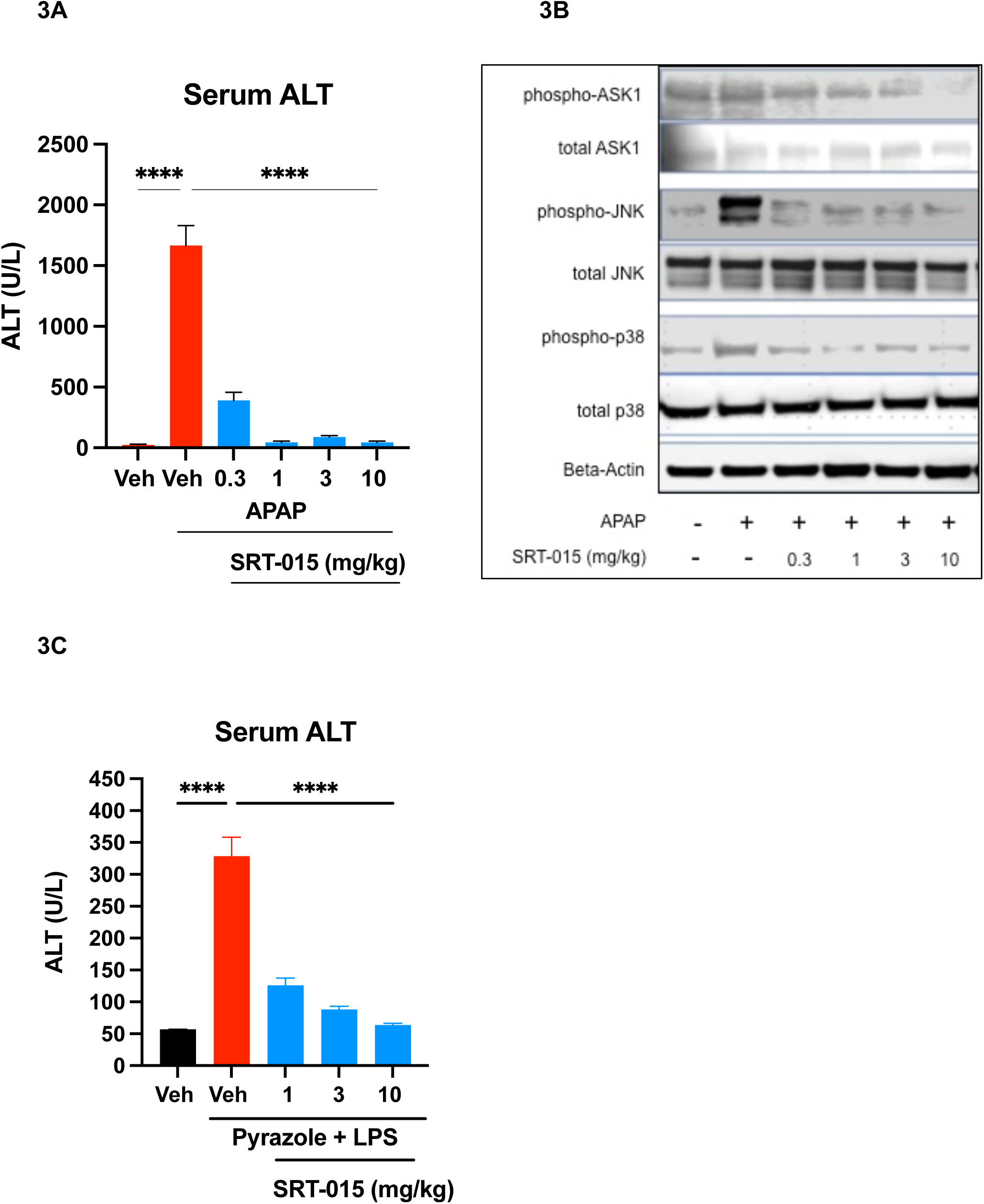
SRT-015 In Vivo Efficacy and Target Engagement in Acute Liver Injury Models. (A) SRT-015 treatment effect on serum ALT 6h after APAP administration in mice (mean ± SEM; n=6). ANOVA and Dunnett’s multiple comparisons test vs APAP administration alone. ****P<0.0001. (B) Western blot analysis after therapeutic dosing of SRT-015 on JNK, p-JNK, p38 and p-p38, ASK1 and p-ASK1 in liver samples collected 6 h post APAP administration. (C) SRT-015 treatment effect on plasma ALT evaluated 24 h after Pyrazole + LPS treatment (mean ± SEM; n = 6-13). Statistical analysis by ANOVA followed by Dunnett’s multiple comparisons test vs Pyrazole + LPS treatment. **** P< 0.0001.

### Acute Pyrazole LPS model of ALD

In this acute ALD model, pyrazole was administered for 2-days to induce CYP2E1 (simulating the alcohol effect) prior to SRT-015 and LPS administration. LPS administration emulates inflammation caused by gut bacteria that penetrate the intestinal wall after binge drinking. Serum ALT was evaluated as a marker of liver injury 24 h after LPS administration and SRT-015 treatment at 1,3 and 10 mg/kg significantly (P<0.001) decreased ALT levels compared to the Pyrazole + LPS group (Fig. 3C). In this model, the MED of SRT-015 was 1 mg/kg.

### Therapeutic DIO-MASH mouse model

After 38 weeks of DIO-MASH diet, mice with biopsy-confirmed fibrosis and steatosis were randomized and treated with SRT-015 or selonsertib for 12 weeks, as shown in Fig. 4A. At 50 weeks, lean chow, DIO-MASH control, SRT-015-treated and selonsertib-treated DIO-MASH mice were evaluated using clinical measures and histological, biochemical, and RNAseq methods.

**Fig. 4.**
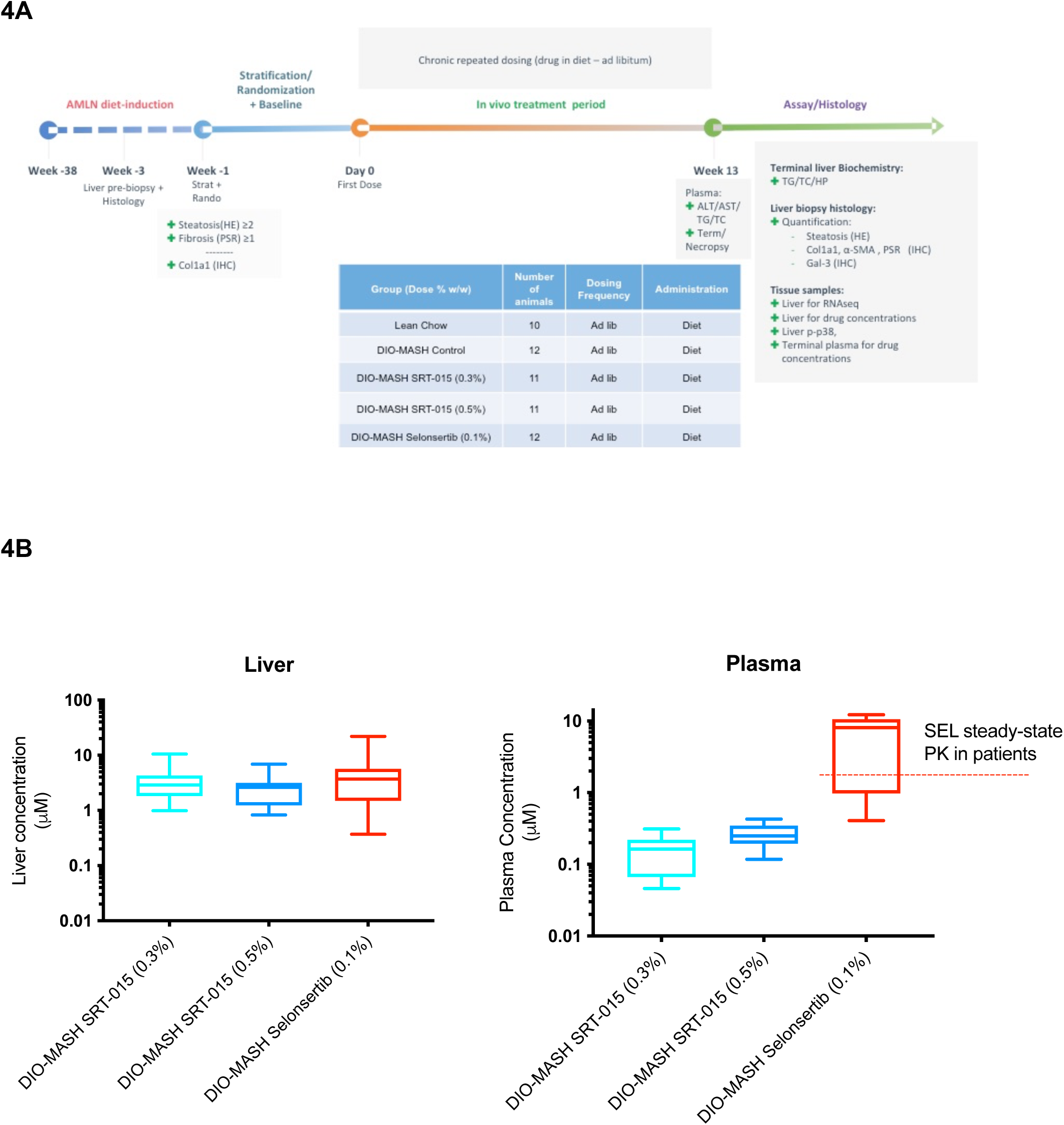
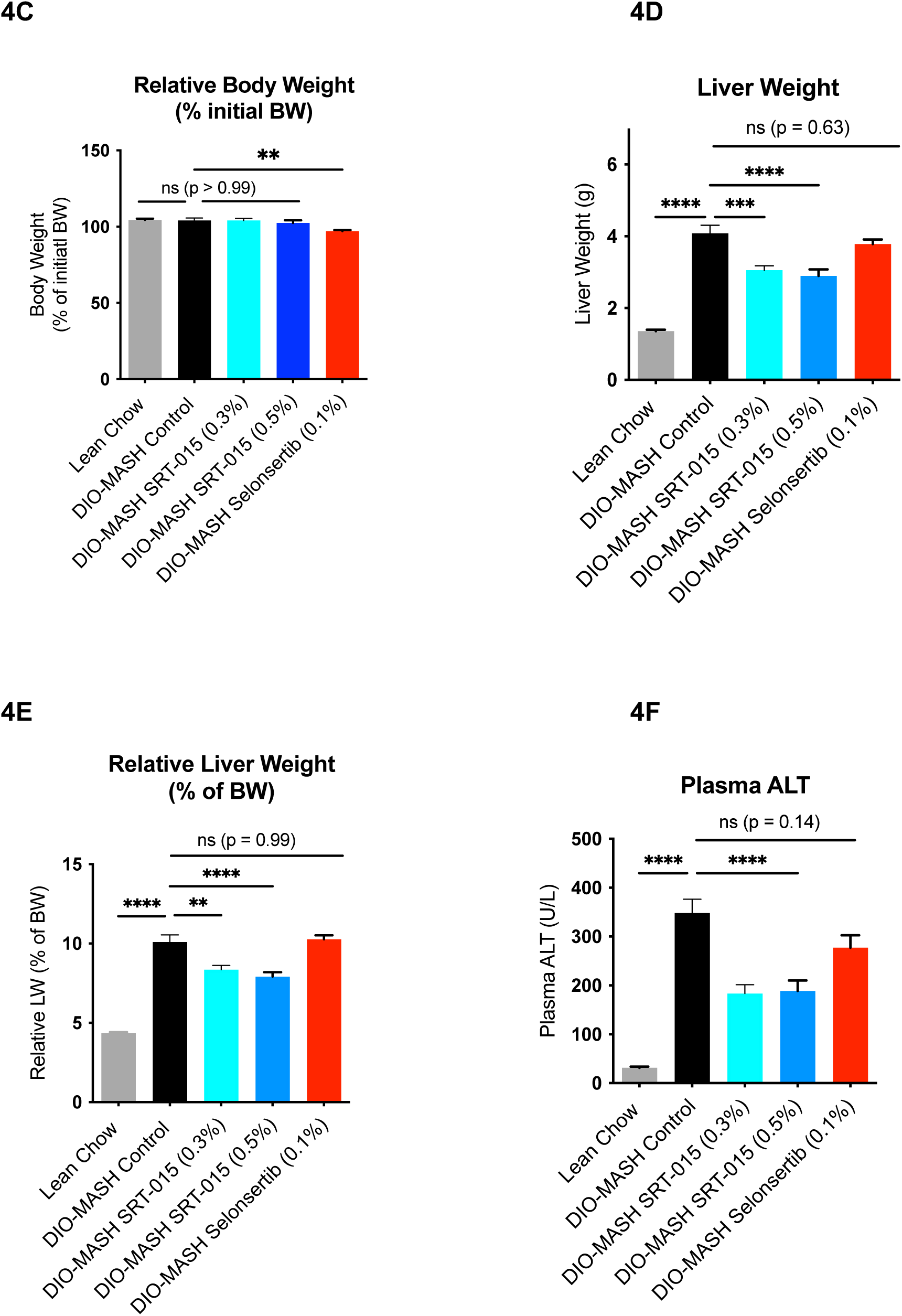
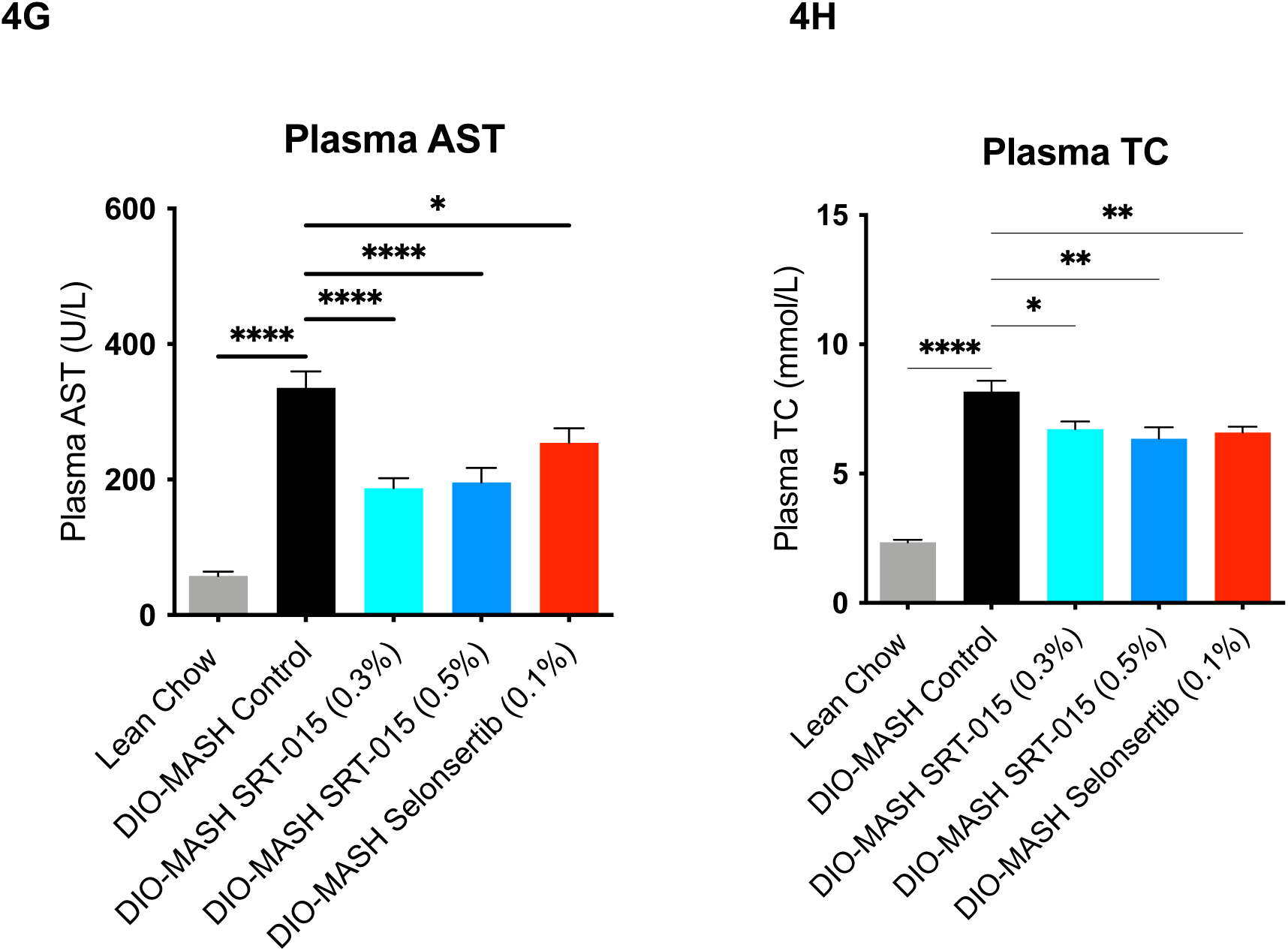
Study design, terminal drug concentrations, BW and LW, and plasma liver enzymes in DIO-MASH study. (A) DIO-MASH study design. (B) Drug liver concentrations from SRT-015 or selonsertib fed DIO-MASH mice were well matched. Selonsertib plasma concentrations match the steady-state PK levels observed in patients (C-G) SRT-015 and selonsertib treatment effects BW, liver weight, and plasma ALT, AST and TC levels. All values expressed as mean ± SEM (n=10-12). Statistical analysis by ANOVA followed by Tukey Multiple Comparisons Test vs DIO-MASH control. * P<0.05, ** P< 0.01, ***P < 0.001, ****P < 0.0001.

Terminal SRT-015 and selonsertib liver drug concentrations were well matched at ∼2 μM (Fig. 4B), and as expected low levels of SRT-015 were observed in the plasma (∼0.2 μM). Selonsertib plasma levels (6.5 ± 1.4μM; n = 12) were higher than the steady-state values (1.3 μM) observed in clinical trials at 18 mg (29).

SRT-015 treatment had no effect (p=0.99) on DIO-induced relative body weight (4C) and significantly decreased (P<0.0001) hepatomegaly (Fig. 4D and E) while selonsertib significantly decreased (P<0.01) relative BW with no effect on hepatomegaly (ns, p = 0.63; Fig. 4C-E). Plasma markers of liver injury, the liver enzymes ALT and AST, were elevated (P<0.0001) in DIO-MASH control mice and significantly decreased (P<0.0001) with SRT-015 treatment demonstrating improved liver function (Fig. 4F,G). In contrast, selonsertib had no or limited effects on ALT (ns, P=0.14) and AST (P<0.05) levels in the DIO-MASH mice. Decreased plasma total triglycerides (TC) were observed following SRT-015 (P<0.01) and selonsertib (P<0.01) treatment (Fig. 4H).

The antifibrotic MOA was observed with SRT-015 treatment decreasing all four biomarker assessments of liver fibrosis (Fig. 5A) compared to DIO-MASH control mice. Morphometry assessments demonstrated a significant reduction (P<0.05) in Col1α1, the main collagen in fibrosis, and PSR, a collagen fibrosis marker. Representative Col1α1 and PSR images are shown in Fig. 5B and sFig. 2, respectively. SRT-015 treatment also non-significantly decreased the activated stellate cell biomarker α-SMA. The collagen content in the liver, measured using a conventional HP biochemical method was also significantly (P<0.05) decreased with SRT-015 treatment. In contrast, selonsertib treatment did not decrease liver Col1α1 (P>0.99), PSR (P=0.58) or HP levels (P=0.73) and significantly (P<0.01) increased α-SMA levels.

**Fig. 5.**
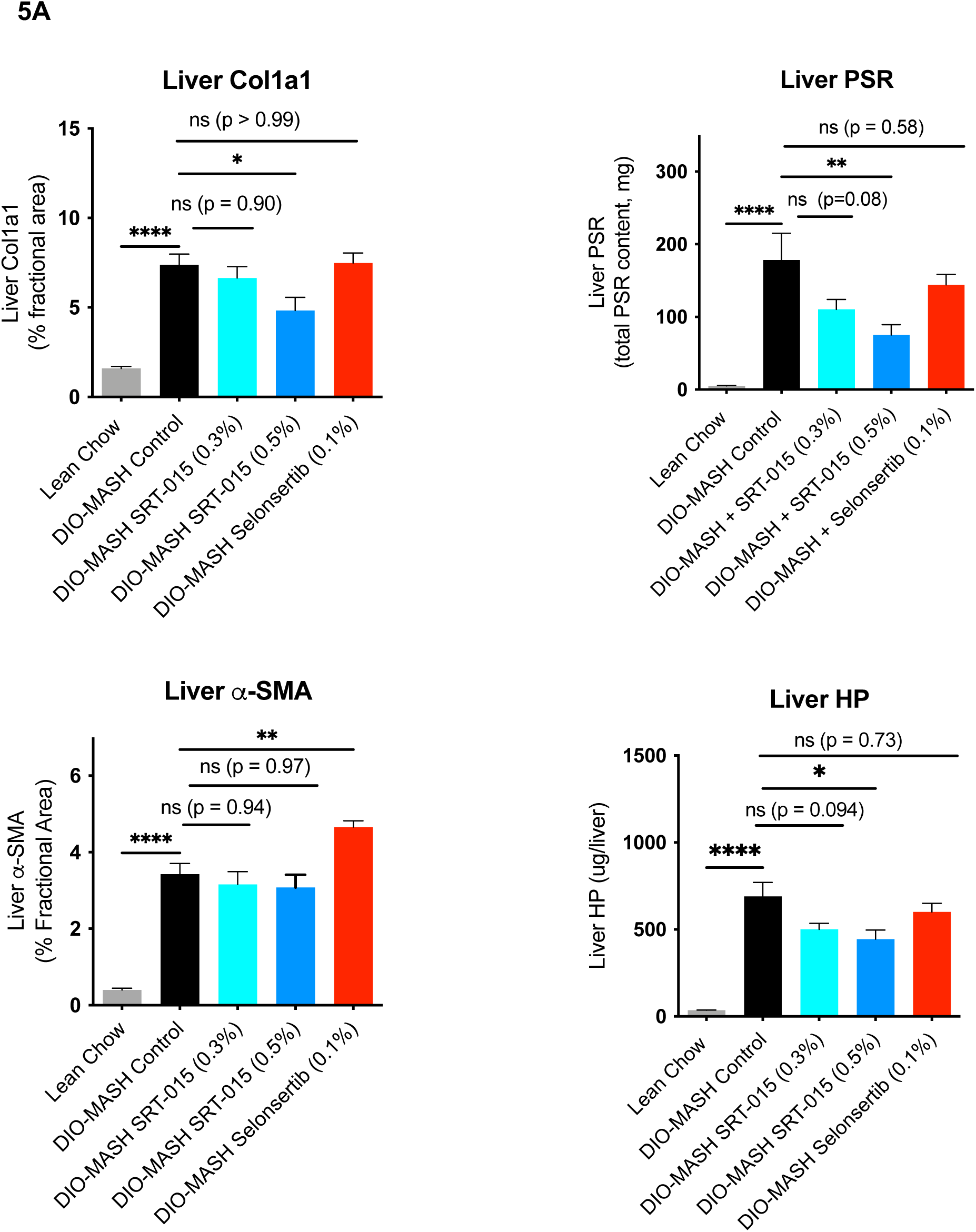
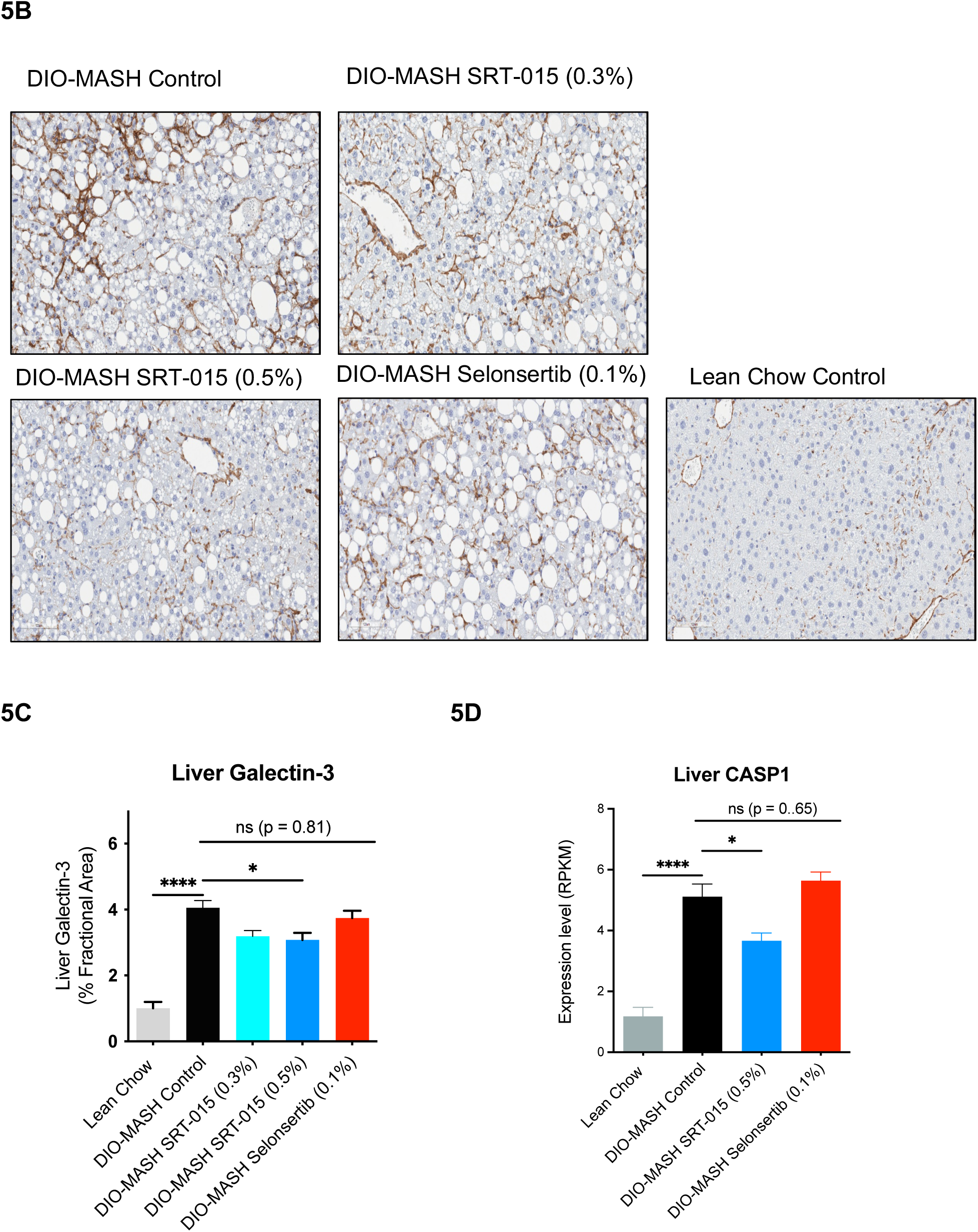
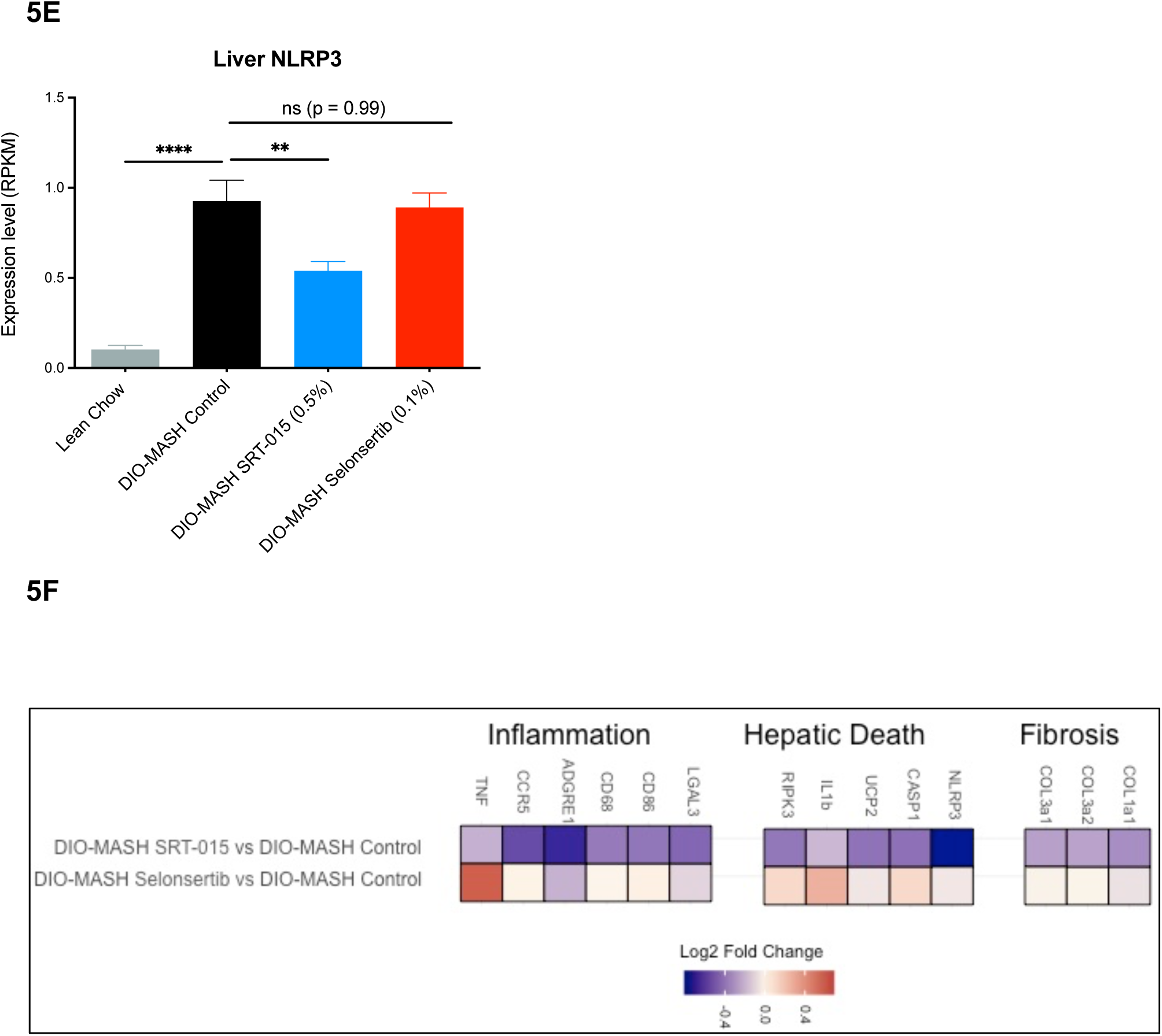
In vivo SRT-015 antifibrotic, anti-inflammatory and anti-apoptotic MOAs demonstrated in the DIO-MASH study (A) DIO-MASH SRT-015 treatment significantly decreased fibrosis assessments. α−SMA was significantly increased by selonsertib treatment (P< 0.01). (B) Representative images of liver stained with anti-Col1α1 (magnification 20x). (C) SRT-015 and selonsertib treatment effects on inflammatory biomarker galectin-3. (D) SRT-015 treatment showing anti-apoptotic MOA by decreased caspace1 (P<0.05) (E) and inhibiting the NLRP3 inflammasome pathway (P< 0.01) (F). Liver RNAseq heat map showing SRT-015 and selonsertib effects on inflammation, hepatic death and fibrosis compared to DIO-MASH controls. All values are expressed as mean ± SEM (n=10-12). Statistical analysis by ANOVA followed by Tukey’s multiple comparisons test vs. DIO-MASH control. * P<0.05, ** P< 0.01, ***P < 0.001, ****P < 0.0001

The anti-inflammatory MOA was observed with SRT-015 treatment in DIO-MASH mice significantly (P<0.05) decreasing galectin-3, an inflammatory monocyte marker, by morphometric analysis (Fig. 5C; representative images sFig. 3) and confirmed by liver RNAseq (LGAL3; Fig. 5F) analysis P<0.01). Additional inflammatory factors, cytokines and monocyte activation factors were down significantly regulated (P < 0.05) in RNAseq analysis including CCR5, CD68, CD86, ADGRE1 (adhesion G protein-coupled receptor E1) and TNF, indicating a broad anti-inflammatory response following SRT-015 treatment (sFig. 4). SRT-015 inhibition of these monocyte recruitment factors will reduce the migration of inflammatory cells into the liver (39).

The apoptosis MOA of SRT-015 was demonstrated by significant decreases in components of hepatic cell death pathways including caspase1 (P<0.05, Fig. 5D) and the NLRP3 inflammasome (P<0.01, Fig. 5E) compared to DIO-MASH control mice. Using RNAseq analysis SRT-015 treatment inhibited other components of the inflammasome/pyroptosis pathway including UCP2 and the necroptosis (RIP3K) pathway (sFig. 5) and confirmed and extended SRT-015 multiple MOAs in the DIO-MASH therapeutic model (Fig. 5F). Selonsertib treatment had no effect on these key pathological mechanisms (Fig. 5F) even with liver drug exposures like SRT-015. In this therapeutic DIO-MASH study, and in a subsequent confirmation study, the SRT-015 minimum efficacious liver concentration was 1-2 μM. With unchanged BW measurements after SRT-015 treatment, the observed improvements in liver function, hepatomegaly, fibrosis, inflammation, and apoptosis can be attributed to the direct effects of SRT-015 treatment in the DIO-MASH mice.

### Target Engagement Phosphorylated p38 (p-p38) IHC DIO-MASH livers

To determine the correlation between p-p38 inhibition and preclinical efficacy, we compared p-p38 IHC using the efficacious compound SRT-015 with the non-efficacious selonsertib in specific liver cell types in the DIO-MASH livers. IHC for p-p38 using liver sections from the DIO-MASH study identified p-p38 positive hepatocytes (large cells) and p-p38 positive small non-hepatocytes (immune/stellate cells; Fig. 6A). The entire section was quantified using segmentation algorithms. DIO-MASH control livers had significantly increased (P<0.01) number of p-p38 positive non-hepatocyte cells (immune/stellate cells; Fig. 6B), but not hepatocytes (Fig. 6C). SRT-015 treatment significantly decreased (P<0.05) the p-p38 positive non-hepatocyte population, demonstrating target engagement and correlating with the observed decrease in fibrosis and inflammation. Selonsertib (Fig. 6B and C) profoundly (P<0.0001) reduced both hepatocyte and non-hepatocytes p-p38 positive cells compared to the DIO-MASH controls with p-p38 positive cell numbers below those of lean chow controls where ASK1 is expected to remain in an inactive state indicating an ASK1-independent mechanism.

**Fig. 6.**
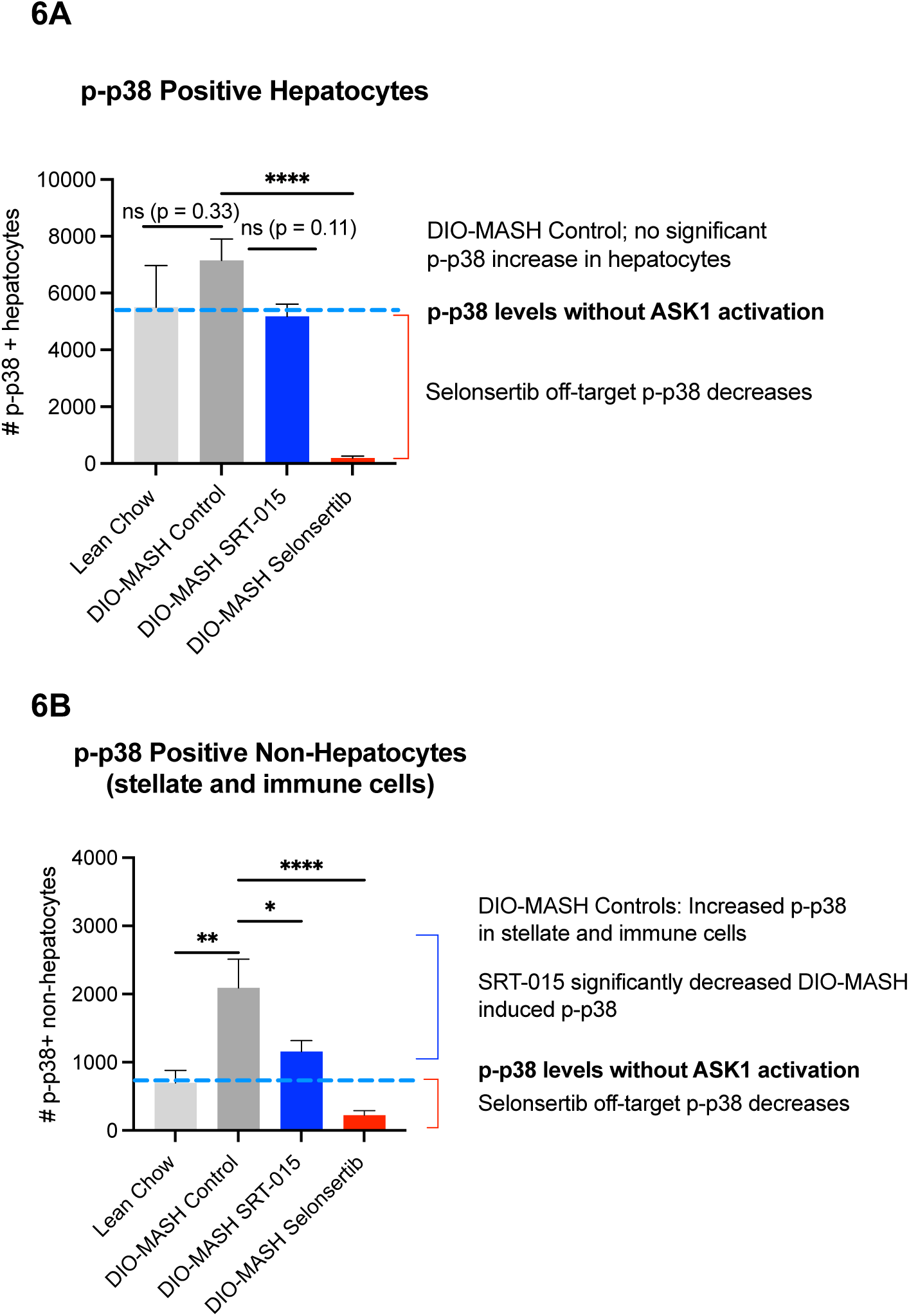
IHC analysis of p-p-38 positive hepatocytes or non-hepatocytes (stellate or immune cells) cell number in DIO-MASH liver study. A) The number of p-p38 hepatocytes in the DIO-MASH controls were insignificantly increased compared to lean chow controls and unchanged with SRT-015 treatment. Selonsertib treatment in DIO-MASH mice decreased p-p38 positive cells below the lean chow controls (red bracket) where ASK1 is not activated indicating an ASK1 independent effect. Dashed blue line indicates lean control p-p38 cell number. (A) In DIO-MASH controls p-p38 positive non-hepatocytes were significantly increased (P< 0.01) compared to lean chow control. SRT-015 treatment significantly decreased p-p38 positive stellate and immune cells. Selonsertib significantly reduced p-p38 cell number compared to DIO-MASH controls levels and also reduced the p-p38 cell number below lean chow controls levels (red bracket) where ASK1 is not activated indicating a non-ASK1 effect. All values expressed as mean ± SEM (n=10-22). Statistical analysis by ANOVA followed by Tukey’s multiple comparisons test vs. DIO-MASH Controls. * P<0.05, ** P< 0.01, ***P < 0.001, ****P < 0.0001

### BDL rat model of cholestatic disease

SRT-015 treatment significantly and dose-dependently decreased BDL-induced fibrosis as demonstrated by decreased PSR staining and HP analysis (Fig. 7 A and B) in this severe model. In Fig. 7C αSMA, a marker of activated stellate cells, was significantly increased by BDL treatment. SRT-015 treatment significantly (P<0.01) and dose-dependently inhibited BDL-induced stellate cell activation. The BDL-elevated blood chemistries were unchanged with SRT-015 treatment. The BW of all BDL groups was decreased, and liver weight increased, compared to sham group and unchanged with SRT-015 treatment (data not shown).

**Fig. 7.**
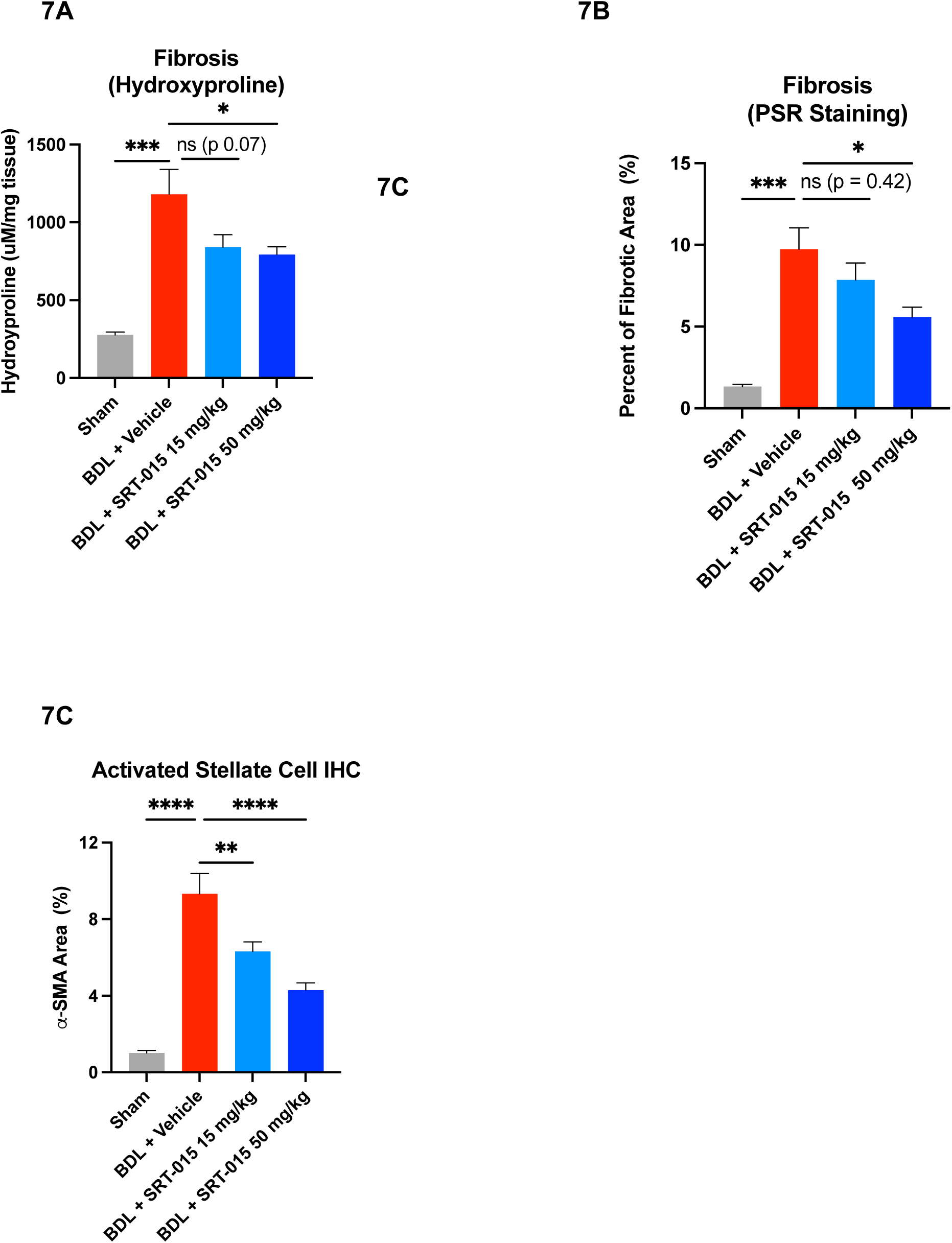
SRT-015 treatment decreased fibrosis and stellate cell activation in BDL rat model. Fibrosis was significantly decreased with SRT-015 daily administration at 50 mg/kg determined by PSR staining (A) or hydroxyproline content (B). Activation of stellate cells was significantly inhibited (C) as shown by decreased α-SMA IHC after 15 and 50 mg/kg SRT-015 treatment. Statistical analysis by ANOVA followed by Dunnett’s multiple comparisons test vs. BDL + vehicle controls. * P<0.05, ** P< 0.01, ***P < 0.001, ****P < 0.0001

## Conclusion

Multiple liver diseases with different etiologies progress due to hepatic cell death, fibrosis and inflammation. A safe and effective therapeutic such as SRT-015, with direct inhibition of multiple liver cell types responsible for these key disease pathologies, would be beneficial for these complex diseases. Here we demonstrated that SRT-015 is a potent, selective and safe inhibitor of ASK1 kinase with direct and pleiotropic MOAs in vitro and in vivo with efficacy in diverse liver diseases including preclinical models of hepatotoxicity, cholestasis, and MASH.

SRT-015 demonstrated a superior cellular functional activity profile comparing with other ASK1 compounds including selonsertib. Other ASK1 inhibitors had off-target hERG liability or cellular cytotoxicity in non-ASK1 activated human cells (no stress inducer present). With SRT-015 treatment, no hERG liability was observed and all required MOAs including anti-inflammatory, antifibrotic, and anti-apoptosis effects on human cells were shown. Further, no off-target cytotoxic effects were observed with SRT-015 treatment in non-activated cells over three population doublings. In contrast, selonsertib and other ASK1 inhibitors exhibited cytotoxicity in primary fibroblasts and hepatocytes in non-activated cells and hERG inhibition. Possible mechanisms for the off-target cytotoxicity of selonsertib including microtubule stabilization (29), and cyclin-dependent kinases 6 (CDK6) inhibition (30).

SRT-015 is an orally bioavailable small-molecule inhibitor of ASK1, with a well-characterized distribution pattern across preclinical species. As demonstrated in mice, rats, and non-human primates, SRT-015 shows a rapid and extensive distribution in the liver and kidney with low levels in the peripheral plasma. The time course of the hepatic distribution in mice revealed that the liver-to-plasma ratio of SRT-015 remained constant over time indicating a lack of tissue accumulation. The low risk of hepatic accumulation after multiple administrations was further confirmed in the 12-week treatment DIO-MASH efficacy study. Importantly, the lack of accumulation of SRT-015 in plasma and liver was also demonstrated in 4-week toxicology studies in rats and non-human primates, where SRT-015 was administered in suprapharmacological doses (unpublished data, A Plonowski). Collectively, the liver-preferred distribution pattern of SRT-015 observed in all preclinical species evaluated suggests that SRT-015 could be particularly useful for the treatment of liver disease with pharmacologically relevant exposure within the liver while maintaining low levels in the systemic circulation. In vivo efficacy was observed with SRT-015 treatment in diverse disease models including acute models of hepatoxicity, cholestatic disease and a therapeutic DIO-MASH model. SRT-015 treatment was effective in acetaminophen overdose or pyrazole/LPS models at 6 h and 2 days, respectively. In a two-week BDL model, SRT-015 treatment significantly inhibited both stellate cell activation and fibrosis. Using a chronic therapeutic biopsy-verified DIO-MASH model a direct comparison of SRT-015 treatment to selonsertib treatment was performed over a 12-week treatment period. The LITMUS consortium identified this model as the top-rated translational animal model for human MASH based on proximal analysis (40). In this model, treatment of DIO-MASH mice with SRT-015 inhibited the DIO-induced increases in hepatomegaly, liver enzymes, fibrosis, inflammation and apoptosis with no body weight loss indicating the direct effects of SRT-015. In contrast, at a dose level mirroring human clinical trial plasma level, selonsertib had no effect on DIO-MASH induced inflammation, fibrosis, apoptosis or other parameters consistent with the negative clinical trial results (27). SRT-015 treatment in DIO-MASH mice decreased all evaluated liver fibrosis parameters. Importantly, liver fibrosis severity is the only feature of MASH that independently predicts liver-related morbidity and mortality (41,42).

To investigate the p-p38 decreases observed with no corresponding efficacy in the selonsertib clinical trial biopsies (24), we performed p-p38 IHC for specific liver cell types in the DIO-MASH study. Evaluation of DIO-MASH control liver sections showed that p-p38 positive cells were significantly elevated only in stellate and immune cells and not in hepatocytes compared to lean controls. SRT-015 12-week treatment significantly decreased DIO-induced p-p38 in stellate and immune cells compared to DIO-MASH control levels demonstrating target engagement in the cell types responsible for fibrosis and inflammation. Selonsertib profoundly reduced p-p38 with no efficacy in this DIO-MASH study in agreement with the clinical trial observations. In addition, selonsertib treatment, uniformly inhibited p-p38 in all cell types below lean chow control levels where ASK1 is expected to remain in an inactive state indicating an ASK1-independent mechanism.

Other off-target liabilities of selonsertib have been well-documented and may have been considered to limit the clinical dose to 18 mg. Selonsertib off-target effects include QT interval prolongation (26) at higher doses (100 mg), that selonsertib is a substrate for CYP3A4 (27) resulting in significant accumulation of a CYP3A4 generated metabolite in preclinical studies and humans (28), and off-target cytotoxicity binding to microtubules or CDK6 (29,30). Additionally, we and others (43) did not observe in vivo efficacy with selonsertib in the LITMUS consortium top-rated translational DIO-MASH model, which demonstrates efficacy with other clinical entities (36,43) including SRT-015. For heterogeneous liver diseases such as MASH, it has been hypothesized that the simultaneous action of different pathways could exert an additive or synergistic effect and optimize the therapeutic results. SRT-015 exhibits multiple direct mechanisms of action in the liver and due to its safety profile could also be used in combination with agents targeting other mechanisms (39). As SRT-015 had no effect on body weight but directly and significantly inhibited fibrosis, inflammation, and apoptosis, combination studies with metabolic agents such as GLP-1 agonists or the recently approved THR-β agonist resmetirom (44) may be beneficial in decreasing fibrosis earlier and to a greater extent.

These preclinical data demonstrate that novel SRT-015 is a specific and potent ASK1 inhibitor with pleiotropic, direct MOAs in vitro and in vivo, with efficacy in diverse liver disease models and support the use of SRT-015 as a potential therapeutic agent in acute and chronic human liver diseases.

## Author contributions

KE and AP: Conceptualization, analysis and interpretation of data, and drafting and editing of the manuscript. SB: conceptualization, analysis, interpretation of data, and editing of the manuscript. MF: analysis and interpretation of data, drafting and editing of the manuscript. NM: drafting and editing of the manuscript and obtaining funding.

## Impact and implications

Human liver diseases, regardless of etiology, share the common pathologies of apoptosis, inflammation and fibrosis. Our study identified SRT-015, a novel ASK1 inhibitor, as a direct inhibitor of these key pathologies in vitro and in vivo. Here we demonstrated SRT-015 treatment is effective in diverse acute and chronic liver disease models with direct inhibition of specific cell types. These findings support SRT-015 as a potential therapeutic for human liver diseases of any etiology.

## Abbreviations

ALD: alcohol-associated liver disease
ALT: alanine transaminase
AST: aspartate aminotransferase
α-SMA: alpha-smooth muscle actin
ADGRE1: adhesion G protein-coupled receptor E1
AMLN: Amylin Liver NASH
APAP: acetaminophen
ASK1: apoptosis signal-regulating kinase 1
BW: body weight
Col1a1: collagen, type I, alpha 1
DIO: diet-induced obesity
ER: endoplasmic reticulum
IC_50_: half-maximal inhibitory concentration
IHC: immunohistochemistry
hERG: human ether-a-go-go-related gene channel
HCC: hepatocellular carcinoma
HP: hydroxyproline
JNK: c-Jun N-terminal kinase
LPS: lipopolysaccharide
MASH: metabolic-disease associated steatohepatitis
MASLD: metabolic dysfunction-associated steatotic liver disease
MED: minimum efficacious dose
MOA: mechanisms of action
NASH: nonalcoholic steatohepatitis
NLRP3 inflammasome: NLR family pyrin domain containing 3
NMP: N-Methylpyrrolidone
p38: 38 kDa mitogen-activated protein kinase
PBMC: peripheral blood mononuclear cells
PK: pharmacokinetics
PSR: picrosirius red
RIP3K: Receptor-interacting serine/threonine-protein kinase 3
ROS: reactive oxygen species
TC: total cholesterol
TGFβ: transforming growth factor bets
TLR4: Toll-like receptor 4
THR-β: thyroid hormone receptor beta
UCP2: uncoupling protein 2.

## Acknowledgements

Sergei Romanov conducted cell functional assays and Martin Madsen conducted RNAseq analysis.

