## Supplemental files for "Novel apoptosis signal-regulating kinase 1 (ASK1) inhibitor SRT-015: Potential therapeutic for multiple liver diseases"

**sTable 1.** Antibody list

**sTable 2**. Pharmacokinetic Parameters of SRT-015 in Rats and Cynomolgus Monkeys

**sFig. 1.** Representative SRT-015 Dose-Response Curves demonstrating direct anti-apoptosis, anti-inflammation and antifibrotic effects using human cells.

**sFig. 2.** Representative images of PSR-stained livers from the DIO-MASH mouse study (magnification 20x).

**sFig. 3.** Representative images of galectin-3 IHC livers from the DIO-MASH mouse study (magnification 20x).

**sFig. 4.** RNAseq analysis of liver tissue corroborated and extended SRT-015 anti-inflammatory mechanisms of action by down-regulation of additional inflammatory factors, cytokines and monocyte recruitment factors.

**sFig. 5.** Anti-hepatocyte cell death MOA was demonstrated with SRT-015 treatment that significantly decreased the expression of hepatic cell death, inflammasome and necroptosis pathways.

**sFig. 6.** IHC p-p38 staining (DAB) and algorithm mask to identify cell type and p-p38 status in liver sections from the DIO-MASH study.

| Description | Manufacturer | Catalog No. | Use |
|---|---|---|---|
| Anti-alpha smooth muscle Actin antibody [1A4] | Abcam | ab7817 | Antifibrosis assay |
| Rabbit anti-JNK antibody | Cell Signaling Technology | 9252 | Western blots |
| Rabbit anti-phospho-JNK antibody | Cell Signaling Technology | 4668 | Western blots |
| Rabbit anti-p38 antibody | Cell Signaling Technology | 9212 | Western blots |
| Rabbit anti-phospho-p38 antibody | Cell Signaling Technology | 9211 | Western blots |
| Rabbit anti-ASK1 antibody | Cell Signaling Technology | 8662 | Western blots |
| Rabbit anti-phospho-ASK1 antibody | Cell Signaling Technology | 3765 | Western blots |
| Rabbit anti-Beta-Actin | Cell Signaling | 4970 | Western blots |
| HRP-linked anti-rabbit IgG (#7074) as secondary antibody | Cell Signaling Technology | 7074 | Western blots |
| Rabbit p-p38 antibody | Cell Signaling Technology | 4511S | DIO-MASH IHC |
| Goat anti-type I Collagen (Col1a1) antibody | Southern Biotech | 1310-01 | DIO-MASH IHC |
| Purified anti-mouse/ human Mac-2 (Galectin-3) Antibody | Biolegend | 125402 | DIO-MASH IHC |
| Recombinant monoclonal rabbit anti-α-SMA antibody | AbCam | 124964 | DIO-MASH IHC |

**sTable 1. Antibody list**

**STable 2**

| Parameter | Rat | Monkey |
|---|---|---|
| **PO administration** | | |
| $C_{max}$ (ng/mL) | 81.4 | 53.4 |
| $AUC_{0\text{-}last}$ (ng*h/mL) | 214 | 348 |
| Liver conc. (ng/g) | 661 (t=1hr) | 1,312 (t=1 hr) |
| (%) Peripheral Bioavailability | 14% | 8% |

**sTable 2. Pharmacokinetic Parameters of SRT-015 in Rats and Cynomolgus Monkeys**
Preclinical pharmacokinetic parameters were determined in plasma samples obtained serially after a single oral (PO, 5 mg/kg) administration of SRT-015 in male Wistar rats and male cynomolgus monkeys.

**sFig. 1. Representative SRT-015 Dose-Response Curves.**

**1A**

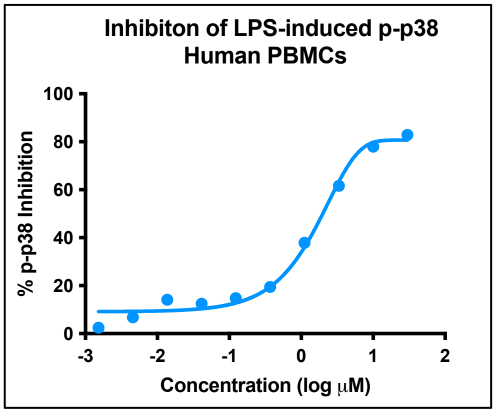


**1B**

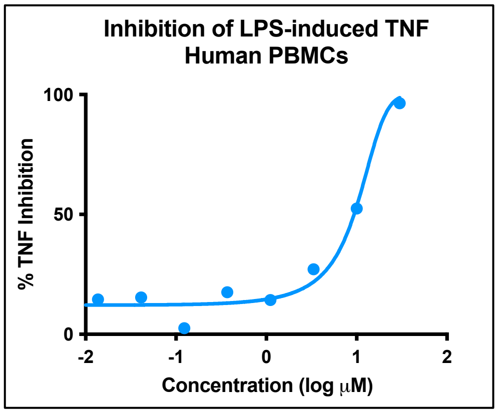


**1C**

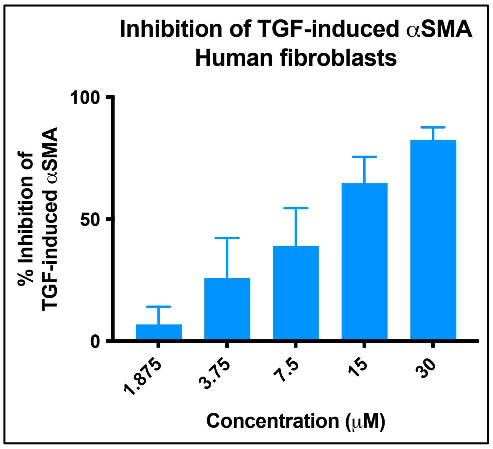


**1D**

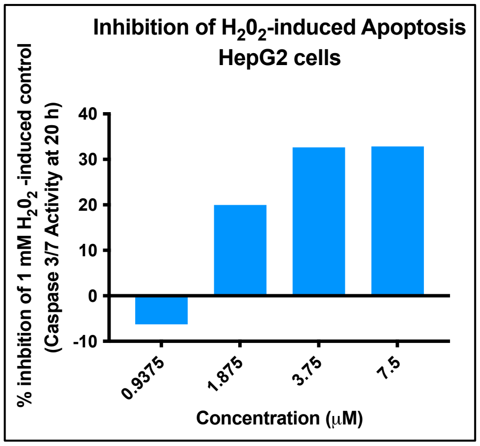


**1E**

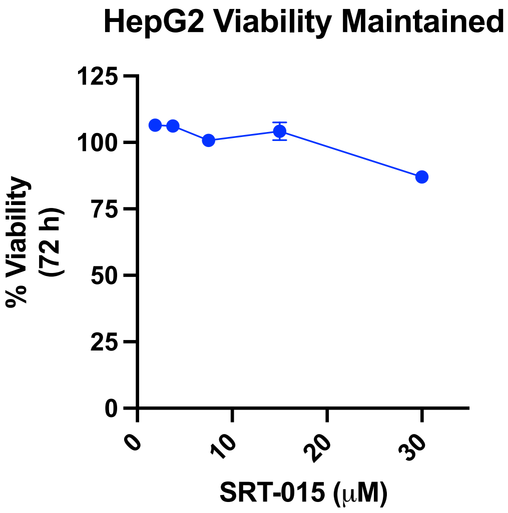

**sFig. 1. Representative SRT-015 Dose-Response Curves Demonstrating Direct, Dose-dependent Inhibition of ROS-ASK1 induced Inflammation, Fibrosis, and Apoptosis, with Target Engagement of the ASK1 Pathway and Without Off-target Cytotoxicity.**

(A) To evaluate the TLR4-ASK1-p38-TNF inflammation pathway, SRT-015 was added 1 h prior to LPS (100 ng/ml) challenge and tested in a 10pt dose response format (0.0015 – 30μM in 0.3% DMSO). LPS challenge activated ASK1 and treatment with SRT-015 dose-dependently decreased p-p38 levels at 20 min, demonstrating target engagement and (B) at 6 h dose-dependently decreased TNF levels demonstrating the anti-inflammatory MOA of SRT-015. (C) The antifibrotic MOA of SRT-015 was demonstrated in human primary fibroblasts tested in a 5pt dose response format. After 1 h preincubation with SRT-015, 0.3ng/mL of TGF was added to activate ASK1 and induce differentiation of fibroblasts into myofibroblasts. After 48 h incubation, α-SMA, a marker of activated myofibroblasts, was dose-dependently decreased with SRT-015 treatment. (D) HepG2 cells after 1h pre-incubation with SRT-015 were stimulated with 1mM $H_2O_2$ for 20h to activate ASK1 and induce apoptosis (Caspase3/7), and cell viability determined in parallel. Caspase 3/7 activity was normalized to cell viability at each concentration point and the percent decreased apoptosis values were used only when cells exhibited 95% or higher viability. Apoptosis was dose-dependently decreased with SRT-015 treatment. (E) No cytotoxic effects were observed with SRT-015 in unstimulated HepG2 cells (no stress protocol) using cells allowed to undergo three population doublings. Comparisons with other ASK1 inhibitors and standards using the cellular assays described here are shown in Table 1.

**sFig. 2**

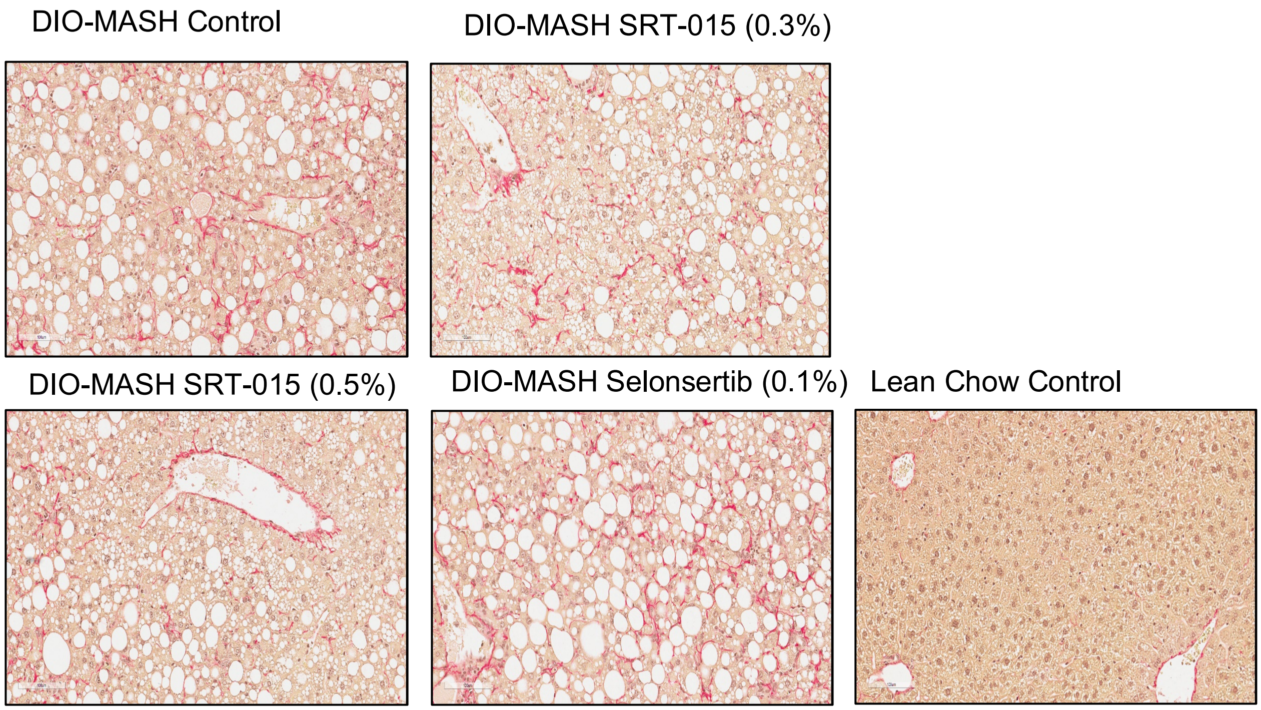


**sFig. 2**

Representative images of Picro Sirius Red (PSR)-stained livers from the DIO-MASH study (magnification 20x).

**sFig. 3**

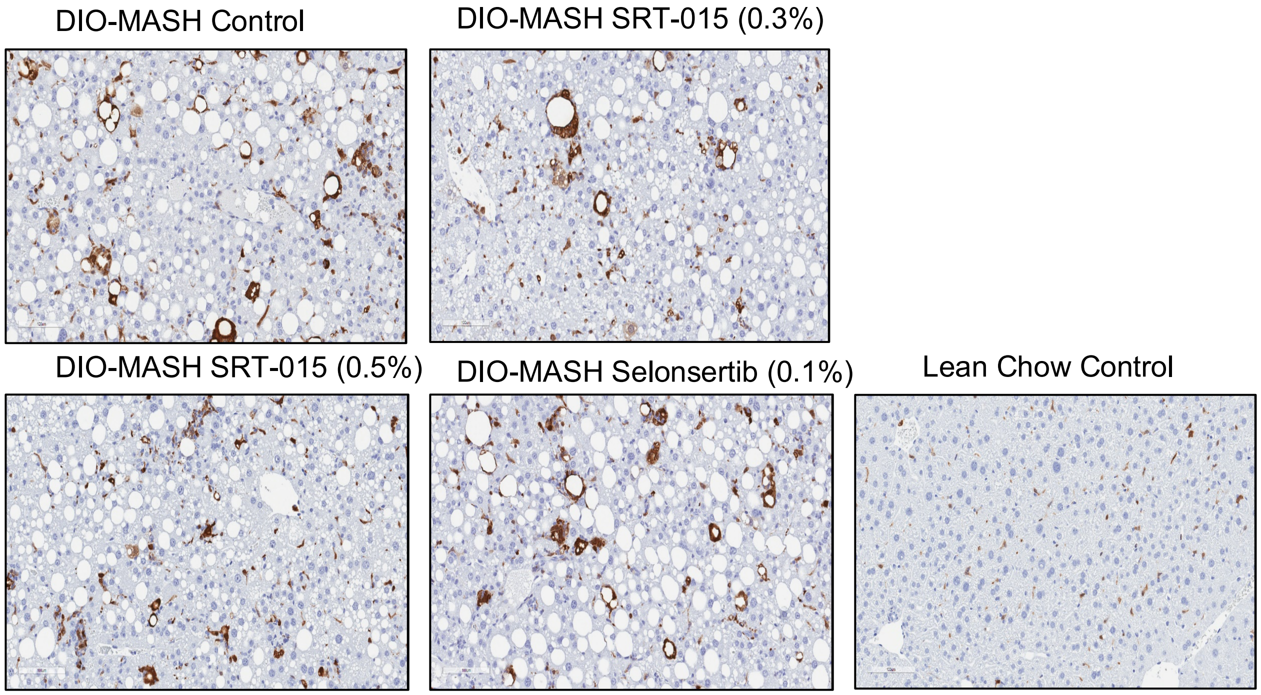


**sFig. 3.** Representative images of galectin-3 IHC livers from the DIO-MASH study (magnification 20x).

### sFig. 4

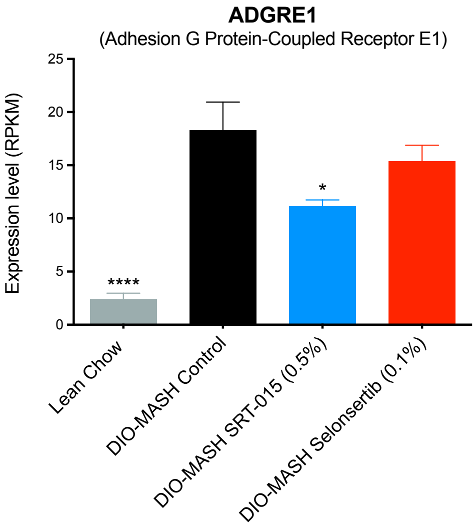


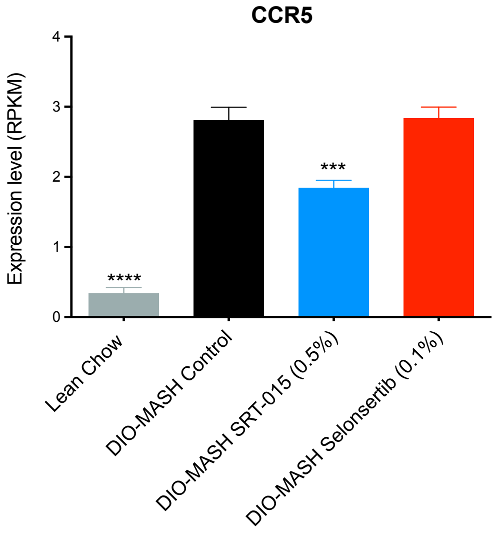


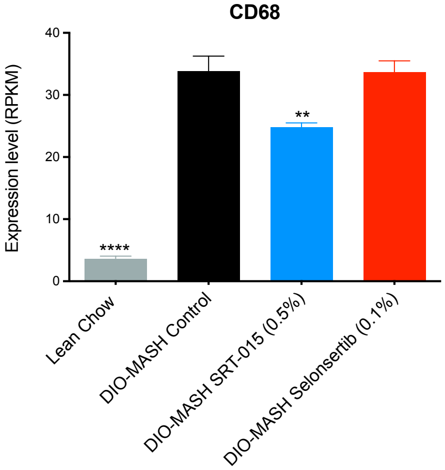


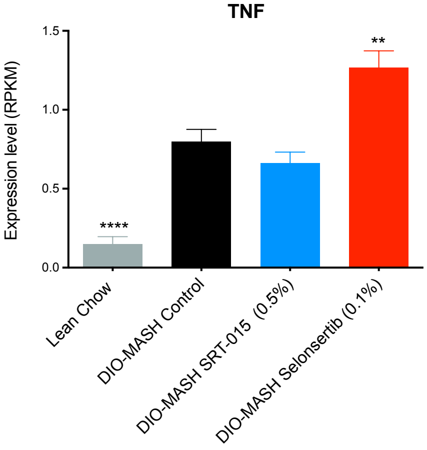


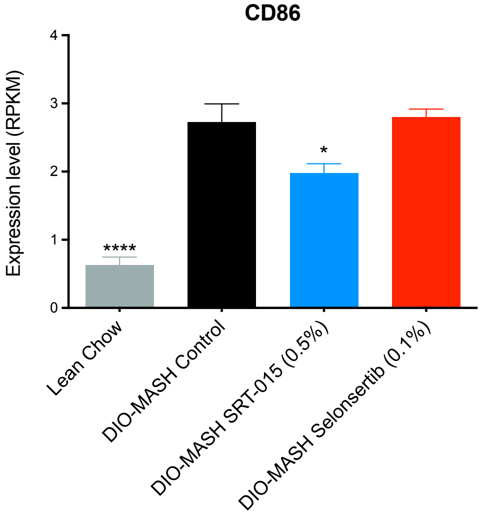


**sFig. 4.** RNAseq analysis of liver tissue corroborated and extended SRT-015 anti-inflammatory mechanism of action by the down-regulation of additional inflammatory factors, cytokines and monocyte recruitment factors. Selonsertib had no significant effect except for an increase in TNF expression. All values are expressed as mean ± SEM (n=10-12). Statistical analysis by ANOVA followed by Tukey's multiple comparisons test; * P<0.05, ** P< 0.01, ***P < 0.001, ****P < 0.0001 vs. DIO-MASH Control

**sFig. 5**

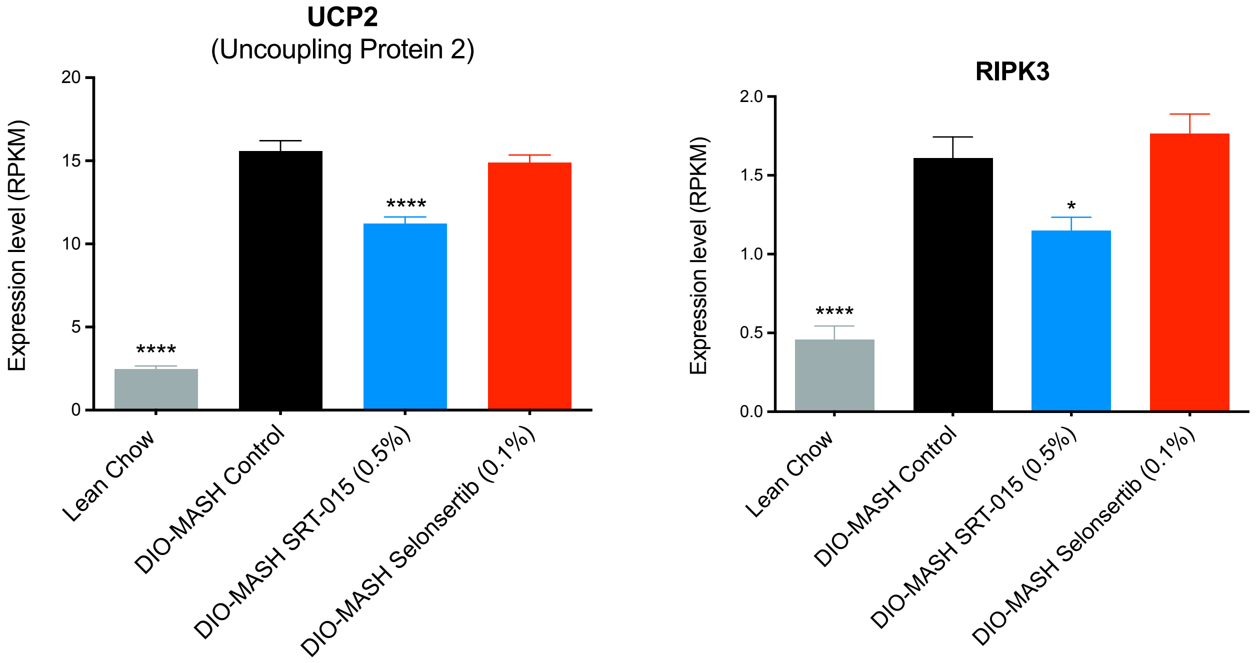


**sFig. 5.** SRT-015 treatment significantly decreased the expression of other hepatic cell death pathways components including UCP2 (inflammasome) and RIPK3 (necroptosis), in DIO-MASH mice. Selonsertib had no effect.

All values are expressed as mean ± SEM (n=10-12). Statistical analysis by ANOVA followed by Tukey's multiple comparisons test; * $P<0.05$, ** $P< 0.01$, ***$P < 0.001$, ****$P < 0.0001$ vs. DIO-MASH Control.

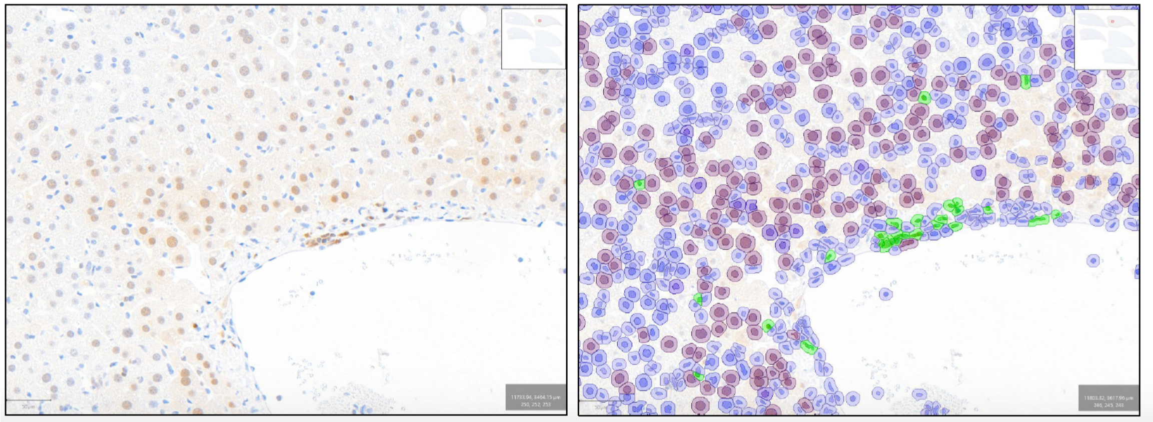

**sFig. 6. IHC p-p38 staining (DAB) and algorithm mask to identify cell type and p-p38 cell number in DIO-MASH liver sections.** A) Left panel shows representative p-p38 staining (DAB) in all cell types in the liver section; right panel with algorithm mask identifying p-p38 negative cells in blue, large p-p38 positive cells (hepatocytes) in brown and small p-p38 positive cells (stellate and immune cells) in green. The entire liver section from all groups was analyzed to determine the number of p-p38 positive non-hepatocytes or hepatocytes.
